# Opposing implications of co-evolutionary lineages and traits of gut microbiome on human health status

**DOI:** 10.1101/2023.05.30.542569

**Authors:** Hao Li, Junliang He, Jieping Liang, Yiting Liang, Wei Zheng, Qingming Qu, Feng Guo

**Affiliations:** State Key Laboratory of Cellular Stress Biology, School of Life Sciences, Xiamen University, Xiamen, China

**Keywords:** wild primates, genomic database, genome reduction, antibiotic resistance genes, carbohydrate-active enzymes

## Abstract

Little is known about the co-evolutionary history of the human gut microbe and its relevance to host physiology. Here, we constructed a gut prokaryotic genomic database of wild primates (pSGBs) and compared it with the human gut prokaryotic database (hSGBs) to define shared co-evolutionary clusters (SCEC-hSGBs) and co-evolutionary traits of hSGBs. We analyzed the evolutionary trends of specific functions like carbohydrate-active enzymes and antibiotic resistance in hSGBs and uncovered host-jumping events and genome reduction tendencies in SCEC-hSGBs. Intriguingly, the SCEC-hSGBs and the super enrichers of the traits (SUEN-hSGBs), which are putatively partially derived from carnivores, showed opposite implications for host health status. Specifically, SUEN-hSGBs are enriched in various diseases, showing a negative correlation with gut biodiversity and disproportionate contributions to the known health-negative marker taxa and metabolite. Our study provides insight into the origin and adaptability of human gut microbes and references for developing probiotics and microbiome-based host health prediction.

## Introduction

Profound and sophisticated metabolic and immune interactions between the gut microbiota and host are primarily built through their co-evolutionary history.^1–4^ According to Janzen’s definition,^5^ providing evidence for bidirectional co-evolution of the host and microbiome is challenging.^4,6^ The term co-evolution in this context refers solely to the evolutionary history of the gut microbiome within a host lineage, regardless of horizontal transferring events. While various factors influence the taxonomic and functional characteristics of the gut microbiome among phylogenetically related hosts, host phylogeny often fairly mirrors the microbial community structure in many cases.^7,8^ The commonness of this phenomenon, coined phylosymbiosis, suggests that continuous co-evolution of gut microbial community across host species may extensively exist.^9^ However, other factors such as sharing similar living environments and diets among closely related hosts may also drive phylosymbiosis.^4,10^

In addition to the community-level co-evolution, a few studies focused on gut microbial intra-lineage co-evolution among hosts,^11–13^ providing insights into long-term interactions between symbionts and hosts. Tracing co-evolutionary lineages is crucial to understand the mechanisms behind co-evolution and co-adaptation between hosts and microbes, since this approach helps to limit interference from recently horizontally transferred taxa with unknown origins. Although both intraspecific (i.e., microevolutionary scale) and interspecific (i.e., macroevolutionary scale) gut microbial lineages have been proposed among related mammals, those co-evolutionary clades spanning hosts diversified millions of years ago are more likely intra-genus multi-specific clusters.^11,12^ So far, a comprehensive list of long-term human gut microbial co-evolutionary lineages transmitted from or among the primate ancestors is lacking. To identify this list, we must compile a reference genome database of gut microbes from wild non-human primates (WNHP), as captive primates exhibit significant disruption in their gut microbiome compared to wild individuals.^14,15^ Therefore, although a genome database of gut microbes from various NHPs has been reported,^16^ an updated genome database excluding captives is necessary.

Evolutionary traits represent ancestral inheritance accumulated over time during continuous adaptation, providing survival benefits for offspring in similar environments.^17,18^ However, such traits can also have adverse effects, such as human mismatch diseases (e.g., diabetes, obesity, and cardiovascular disease) partially or wholly caused by mismatches between long-term adaptation and recent rapid environmental changes.^19,20^ Currently, the co-evolutionary traits of the gut microbiome of *Homo sapiens* and their implications with host health are poorly characterized. The optimal reference for profiling such traits should be the gut microbiome of the closest relatives, i.e., other *Homo* species, but lack of sufficiently analyzable fecal samples from extinct *Homo* species prevents this possibility.^21^ Thus, a suboptimal option is to profile the long-term co-evolutionary trait of the human gut microbiome by referring to our primate cousins.

We previously discovered a multi-species *Prevotella copri*-containing lineage co-evolved with WNHPs and humans.^11^ Here, we aimed to further characterize long-term co-evolutionary human gut microbial lineages and traits and investigated their implications on host health. We built an updated genomic database of gut prokaryotic species containing over 1,600 species from various species of WNHPs. By comparing this with human gut microbial genomes,^22^ we then defined the co-evolutionary species and enriched traits in the human gut microbiome. Interestingly, we found that the co-evolutionary lineages and super-enrichers of evolutionary traits were oppositely correlated with human health status. Our results provided novel insights into how gut microbiota adapts to continuously evolving human niches from an evolutionary perspective. Additionally, we identified previously unidentified gut microbiome biomarkers of health status.

## Results

### An updated gut prokaryotic genome database from WNHPs containing 1,654 species-level genome bins (SGBs)

We collected 346 fecal metagenomes from 25 primate species, including 15 *Macaca thibetana* data contributed by this study, for recovering metagenome-assembled genomes (MAGs) (Figure 1A and Table S1). Therein, 284 metagenomes were derived from WNHPs,^17,23–29^ while the remaining 62 metagenomes were obtained from captive *M. mulatta* (*n* = 6), *M. leonina* (*n* = 4), *M. thibetana* (*n* = 3), *Pan troglodyte*s (*n* = 18), *Gorilla gorilla* (*n* = 22), *Lemur catta* (*n* = 5), and *Papio anubis* (*n* = 4).^23,28,30^

**Figure 1.**
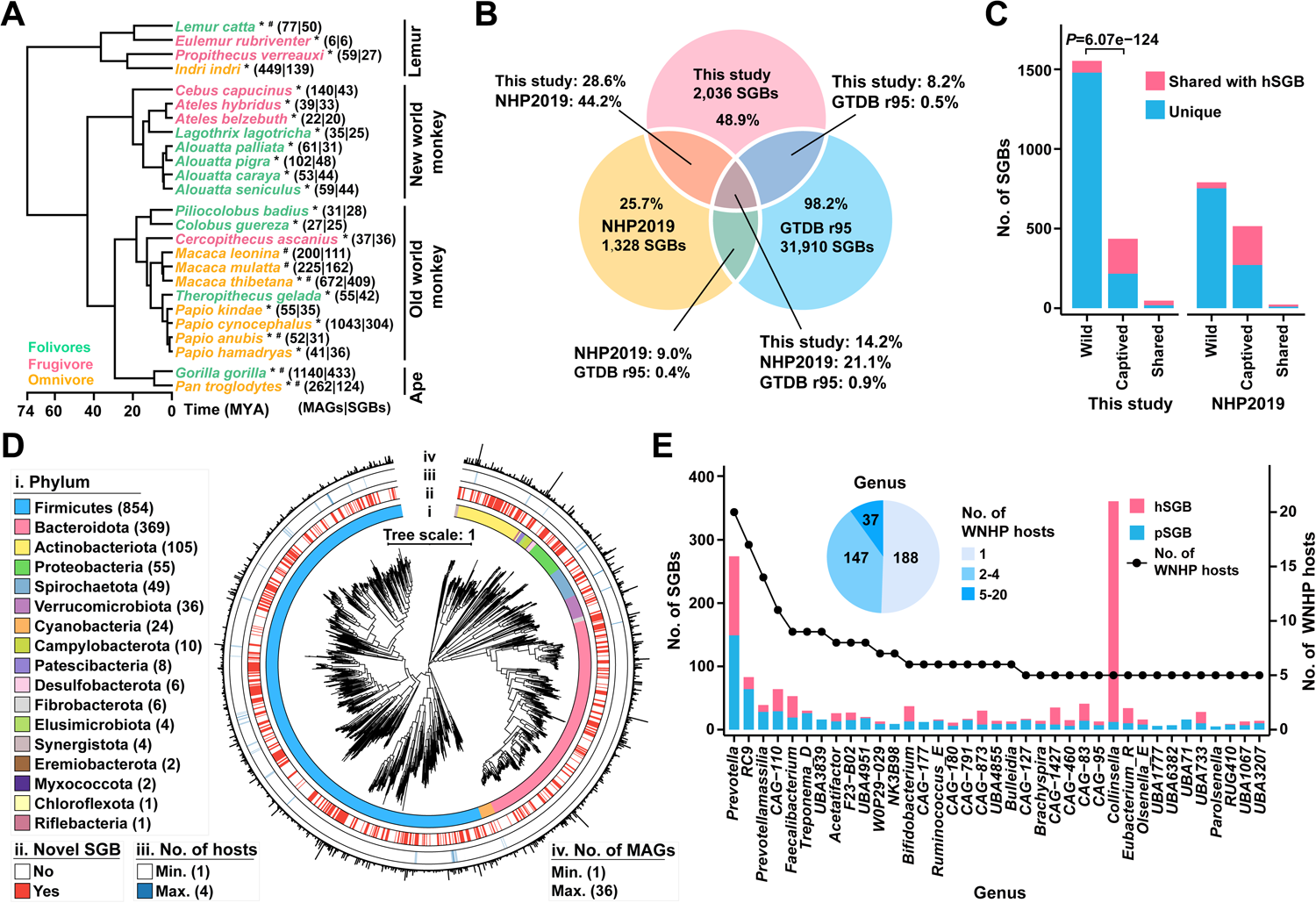
The updated genomic database of gut microbes from WNHPs. **A)** Phylogenetic tree of 25 NHPs based on the evolutionary timescale. Hosts from the wild and captived sources are labeled by * and #, respectively. **B)** Venn diagram of shared SGBs (ANI >95%) between the NHP database of this study, the NHP database from NHP2019, and GTDB r95. **C)** Shared SGBs between hSGBs and our NHP database or NHP2019 (ANI >95%). **D)** Phylogenetic tree of pSGBs based on concatenated 120 universal single-copy genes. Only 1,536 bacterial SGBs with gaps in <60% of alignment columns are shown. **E)** Pie chat indicates the host number of non-singleton genera (≥2 SGBs) of pSGBs, and the bar plot depicts the genome number of pSGBs and hSGBs in genera with ≥5 WNHP hosts.

In total, we retrieved 4,942 bacterial and archaeal MAGs (>50% completeness and <5% contamination) representing 2,036 SGBs (clustered by average nucleotide identity (ANI) >95%, see Table S2 for the information of each MAG and SGB) from the metagenomes. These MAGs represent 54.6% (median value) of the corresponding metagenomes, indicating high coverage of the SGBs in the gut microbiome of primates. Nearly half of the SGBs (41.6%) were high-quality genomes with completeness >90% and contamination <5% (Fig S1A). Strikingly, 78% of the SGBs lacked conspecifics in the GTDB-r95 database (Figure 1B),^31^ suggesting that a substantial number of these SGBs may uniquely distribute in the gut of primates. Even compared to the gut microbial genomic database of NHPs constructed by Manara et al. (2019) (NHP2019),^16^ over half of the SGBs (*n* = 1,163) were exclusively detected in our database (Figure 1B).

To evaluate the impact of anthropogenic disturbance on the captive individuals, we sorted the 2,036 SGBs into three catalogs based on their source: wild individuals, captive individuals, and shared by both (Figure 1C). The shared SGBs appeared in a low percentage (*n* = 47). The ANI values were determined between the SGBs in each catalog and the human gut SGB collection (hereafter, hSGBs, *n* = 3,779).^22^ As expected, captive primates harbored a much higher ratio of hSGBs-conspecific SGBs than the wild ones (220 in 436 vs. 74 in 1,553, *P* = 6.07e−124, Chi-Square test). Similar proportions were detected for the NHP2019 (Figure 1C). A plausible inference is that these hSGBs-conspecific SGBs inhabiting in captive primates were horizontally transferred from humans or related environments under captivity. Therefore, to eliminate the effects of captivity, all SGBs solely contributed by captive individuals were excluded. Finally, after combining the remaining 1,576 SGBs and the non-redundant fraction from wild individuals of NHP2019, the final database contained 1,654 SGBs (1,635 Bacteria and 19 Archaea) from 23 WNHP species (hereafter, pSGBs).

The pSGBs are affiliated with 19 phyla and 1,436 of them can be classified into 372 genera according to the GTDB taxonomic system (Figure 1D and Table S3). Here we congregated three phyla (Firmicutes, Firmicutes_A, and Firmicutes_C) into Firmicutes. The top five phyla, namely, Firmicutes, Bacteroidota, Actinobacteriota, Proteobacteria, and Spirochaetota, constitute 91.8% of the total (Figure 1D). Only 2.7% (*n* = 45) SGBs harbored MAGs from multiple WNHP species, with a maximum of four shared hosts. However, it does not imply that the pSGBs from different primate hosts were remotely related. Among the 222 non-singleton genera (containing two or more pSGBs), 184 had SGBs found in multiple host species, and 37 of them can be detected in ≥5 host species (Figure 1E). *Prevotella*, with 149 pSGBs, has the broadest host range (contributed by 20 WNHP species). Furthermore, hSGBs were detected in 234 pSGB-containing genera, encompassing 1,128 pSGBs and 1,988 hSGBs (Table S4). The high proportion of congeneric hSGBs and pSGBs suggested the prevalence of long-term co-evolutionary history between a large number of microbial lineages and primate hosts.

### Distributive feature of carbohydrate-active enzymes (CAZys) and antibiotic resistance genes (ARGs) in hSGBs and pSGBs

We investigated the distribution of two types of functional genes, CAZys (including only glycoside hydrolase (GH) and polysaccharide lyase (PL)) and ARGs, between pSGBs and hSGBs, as selective pressures on these genes likely differ substantially between WNHPs and modern humans. Overall, the number of CAZy families and genes in pSGBs and hSGBs generally decreased from lemurs and monkeys to apes, then to humans (Figure 2A, B). The average number of CAZy genes is 28.5 in hSGBs and 38.0 in pSGBs. The dietary type also impacts the number of CAZy families (Fig S2A). Further analysis showed numerous GH families enriched in pSGBs and relatively few in hSGBs (Figure 2C and Table S5). Among the top enriched GH families in pSGBs, many are related to the degradation of cellulose (GH5_2 and GH5_4), xylan (GH43_18), and arabinose-related glycoside (GH43_18 and GH53), consistent with the higher intake of these plant glycans in wild primate versus human diets. In contrast, enzymes targeting glucose-related and galactose-related glycans (GH1, GH4, GH32, and GH112) were enriched in hSGBs (Figure 2B and Fig S2B).

**Figure 2.**
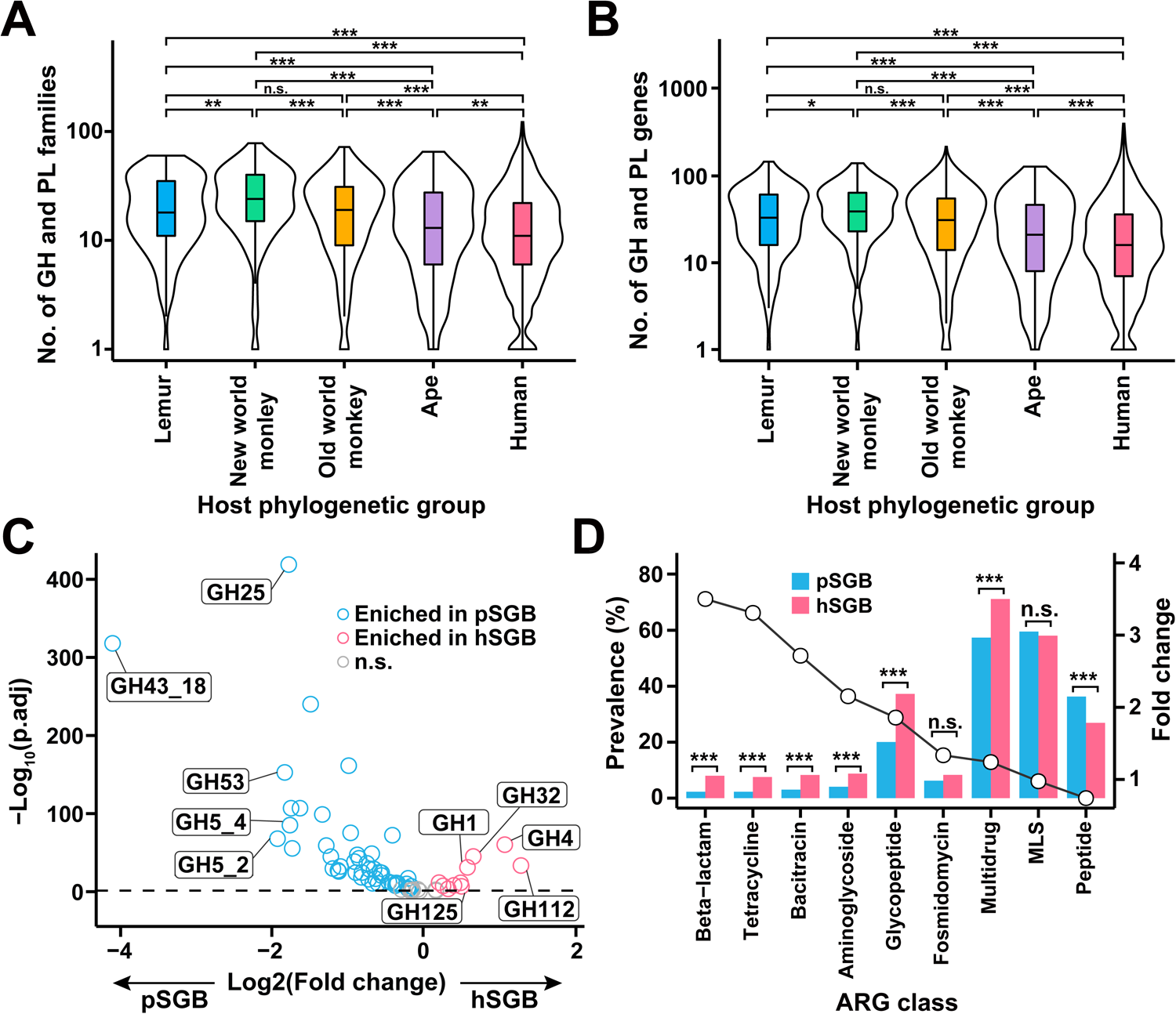
CAZy and ARGs profile of pSGB. **A, B)** Number of CAZy families or genes of SGBs from different primate host phylogenetic groups. Two-tailed Mann-Whitney U-test. **C)** Volcano plot of enriched CAZymes in pSGBs (blue) or hSGBs (red). Only CAZymes with >10% prevalence in either database were shown, and only the top five enriched CAZys of each database were labeled. The dashed line indicates *P*_adj_ = 0.05. Fisher’s exact test. **D)** Most prevalent ARG classes in pSGBs and hSGBs. Only ARG classes with a prevalence >5% in either database were shown. Fisher’s exact test. *, *P*_adj_ < 0.05; **, *P*_adj_ < 0.01; ***, *P*_adj_ < 0.001; n.s., not significant.

For ARGs, the top enriched classes in hSGBs encode resistance to beta-lactam, tetracycline, bacitracin, aminoglycoside, and glycopeptide (ranked by enriching fold of prevalence, Figure 2D and Table S6). Conversely, highly prevalent ARG classes such as macrolide-lincosamide-streptogramin (MLS) and antimicrobial peptides were detected in pSGBs at a similar or even higher rate. These results support the directional selection of the human gut resistome over less than 100 years of clinical antibiotic use.

### Defining shared co-evolutionary SGB clusters

To identify all detectable co-evolutionary SGB clusters, we calculate the ANI values between congeneric pSGBs and hSGBs, as it is a comprehensive measurement of the evolutionary distance between intra-genus genomic pairs.^32^ We hypothesize an operational ANI threshold, under which the generated SGB clusters can optimally represent the diversified offspring in primate hosts from each ancestral bacterial species (i.e., the balance between conservative and radical), even though we cannot confidently determine the speciation time for the ancestor bacteria (i.e., within the primate host or not). By stepwise increasing the ANI threshold, the split ratio of SGB clusters was evaluated for determining the operational threshold (see Methods and a diagram in Figure 3A). We observed the first significant increase in the split ratio for non-singleton SGB clusters when increasing the ANI threshold from 77 to 78% (FDR-corrected *P* = 1.3e−7, Fisher’s exact test, Figure 3B). Over three-quarters (285 in 360) of non-singleton ANI-77% clusters split from ANI-78% to ANI-83% (see Fig S3A, notice that ANI-83% is a widely recognized lower limit of ANI for closely related prokaryotes),^33^ indicating that most ANI-77% clusters comprised remotely related species. Moreover, the corresponding host divergent time for the split-out SGBs also showed a decreasing trend since ANI-77%, especially when excluding hSGBs (Fig S3B). These results corroborate ANI-77% as the optimal operational threshold for defining the co-evolutionary SGB clusters.

**Figure 3.**
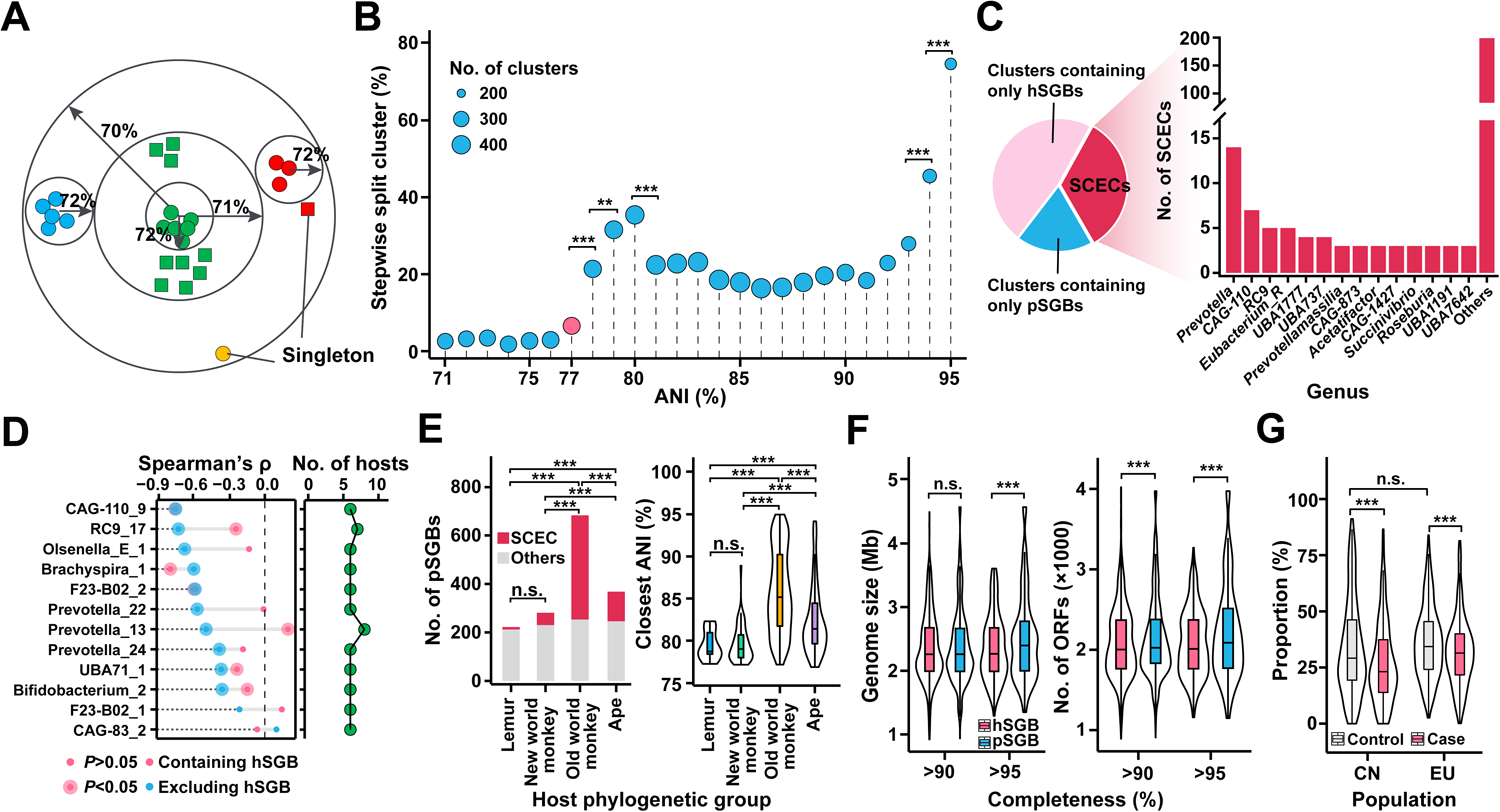
Defining and characterizing SCECs. **A)** Schematic of the definition of split clusters using stepwise increasing ANI values. **B)** Determining the operational threshold of defining SCEC by stepwise increasing ANI values. ANI=77% was selected as the threshold to define the co-evolutionary SGB clusters. Fisher’s exact test. **C)** Pie chat shows the proportion of SCECs in non-singleton clusters under ANI=77%. The bar plot depicts the number of SCECs in the genera. Only genera with SCECs ≥3 are shown. **D)** Correlation between the ANI value of SGBs within the SCEC with the divergence time of their hosts (left panel). Only SCECs with ≥ 6 host species were shown (point, right panel). **E)** The proportion (left panel, Fisher’s exact test) of SCEC-pSGB and their closest ANI to hSGB in the four primate phylogenetic groups (right panel, two-sided Mann-Whitney U-test). **F)** Comparison of genome size and the number of encoding ORFs between pSGB and hSGB within SCEC under different completeness thresholds. Paired two-sided Student’s *t*-test. **G)** The proportion of SCEC-hSGBs in CN and EU populations based on the ICG database. Two-sided Student’s *t*-test.

Under the threshold, we defined 779 co-evolved non-singleton SGB clusters in total (Table S7), of which only 262 contained both hSGB and pSGB (shared co-evolutionary clusters, SCECs). As shown in Figure 3C, *Prevotella* has the most SCECs (*n* = 14), echoing its high species-level diversity in both pSGBs and hSGBs (Figure 1E). Among these SCECs, the *Prevotella*_13 SCEC associates with the largest number of WNHP hosts, showing highly overlapped pattern with the *P. copri*-containing lineage we proposed earlier.^11^ For the 12 SCECs detected in ≥ 6 host species (i.e., ≥ 5 WNHP species and human), we observed overall negative correlations between the ANI value and host divergent time (Figure 3D). Intriguingly, the negative correlations strengthened when excluding hSGBs. Taken together with Fig S3B, this observation suggests that some hSGBs may remotely transfer from other primates, initiating independent evolution in human ancestors within their long-term co-evolutionary history. We also find that old world monkeys have an even higher proportion of SCEC-pSGBs than apes, while both are much higher than lemurs and new world monkeys (Figure 3E). A similar pattern was also observed for their ANI to the closest SCEC-hSGBs (Figure 3E). These results indicate that the remote horizontal transfer events substantially impacted the distribution of SCEC-hSGBs in the gut of modern humans.

Defining SCECs favors profiling the evolutionary trajectory of hSGBs relative to their pSGB counterparts. We compared genome size for each intra-SCEC pair of hSGBs and pSGBs, finding apparent genome reduction in hSGBs (mean 5.8% for genomes >95% completeness, Figure 3F). The reduction appears attributable to both gene loss and changes in gene length but not encoding density (Figure 3F and Fig S3C). Meanwhile, SCEC-hSGBs have smaller genomes than other hSGBs (*P* < 0.001, two-sided Student’s *t*-test, Fig S3D). In terms of functions, such genome reduction can be partially explained by the loss of CAZy genes (Figure 2B). Additionally, the top shrinking gene categories (annotated by Clusters Orthologous Genes, COGs) are those related to cell motility and energy production & conversion (Fig S3E).

We also investigated the distribution of SCEC-hSGBs in two human populations (China and Europe) using the IGC database.^34^ We found SCEC-hSGBs comparably distributed in healthy populations from China and Europe (Figure 3G). Interestingly, we observed a decrease of SCEC-hSGBs in disease individuals for both China (type-2 diabetes, T2D) and Europe (inflammatory bowel disease, IBD, *P* < 0.001, two-sided Mann-Whitney U test, Figure 3G), preliminarily suggesting SCEC-hSGBs may positively implicate with host health.

### Profiling co-evolutionary traits of hSGBs

Given our interest in gained or strengthened functions of hSGB, we focused on the hSGB-enriched co-evolutionary traits (annotated by COG). We identified 839 candidate COGs based on prevalence and abundance by comparing all hSGBs and pSGBs. Among them, as recaptured in metagenomic data, 695 are defined as the co-evolutionary traits of hSGBs compared with pSGBs (Figure 4A and Table S8, see Method and Figure S4A-C). Further examination revealed that only a small fraction (135, 19.4%) of these trait COGs are also enriched in SCEC-hSGBs compared with their SCEC-pSGB counterparts. Nevertheless, the remaining 560 COGs exhibit a strong positive correlation with the 135 COGs among all hSGBs, and SCEC-hSGBs typically had low numbers of total trait COGs (Figure 4A), possibly determined by their small genome size.

**Figure 4.**
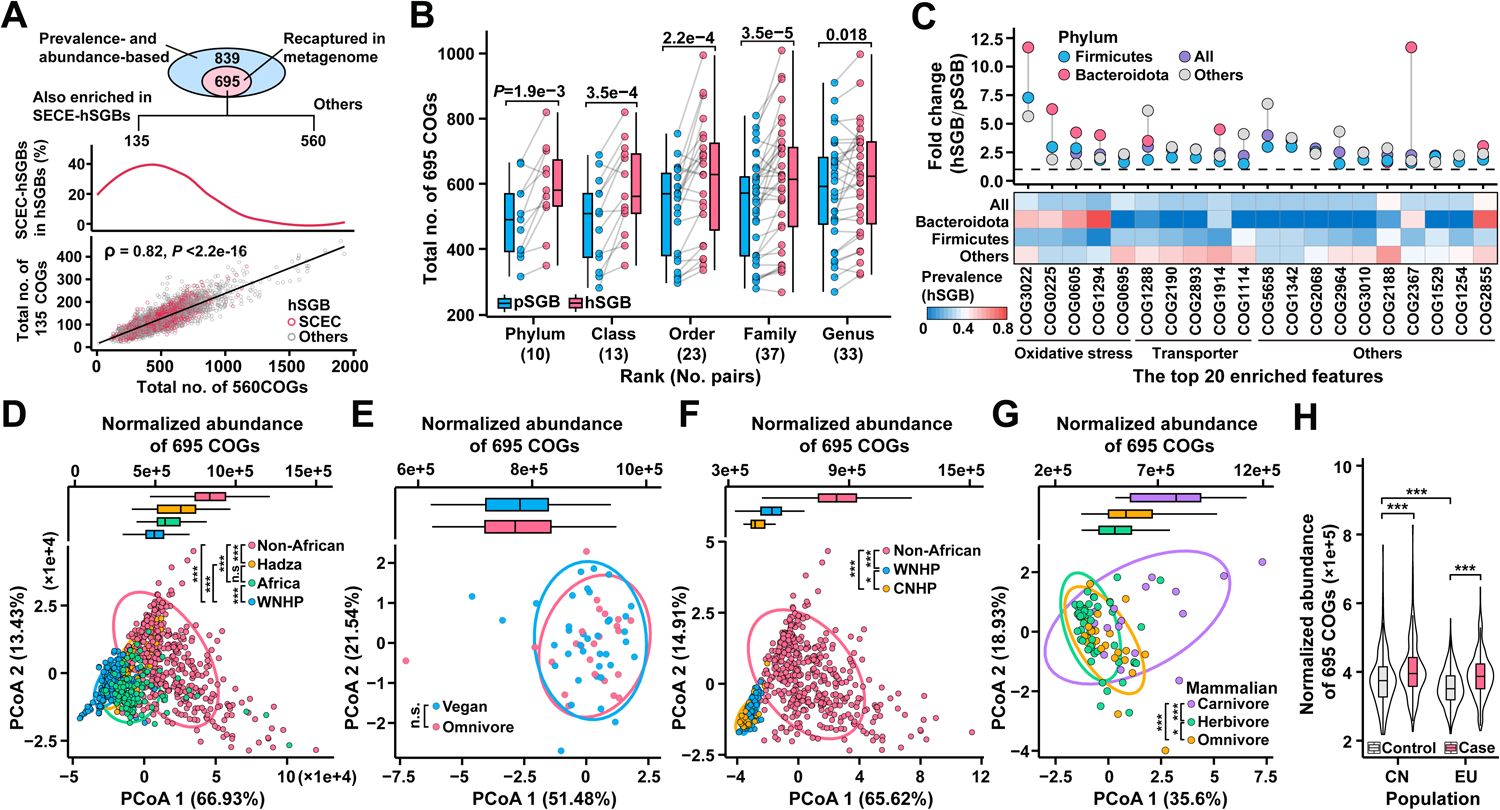
Defining the co-evolutionary traits of hSGBs and constraints affecting their distribution. **A)** Overview of defining the co-evolutionary traits of hSGBs (top panel). The medium panel depicted the distribution of 695 COGs in SCEC-hSGBs. The bottom panel showed the correlation between 560 COGs and 135 COGs in hSGBs. **B)** Comparison of 695 co-evolutionary traits between pSGB and hSGB. Genomes with completeness >90% were considered, and the difference of 695 co-evolutionary traits between pSGB and hSGB is calculated based on the average value of the total copy number within the genome. The number of taxa pairs at each rank is shown in parentheses. **C)** Distribution and functional profile of the top 20 significantly enriched COGs in hSGB (Fisher’s extract test with FDR corrected *P*<0.05). The dropline showed the fold change of the prevalence of COGs in hSGB compared pSGBs at the phylum level. Only COGs with a prevalence >5% in either database were displayed. The heatmap showed the prevalence of these COGs in hSGB at the phylum level. The dashed line represents the prevalence of COG is equal in both databases. **D-G)** Boxplot (top panel) indicated the abundance differences of the 695 COGs across metagenomic groups, and Euclidean distance PCoA based on the relative abundance of 695 COGs (bottom panel) shows its distribution pattern. Ellipses cover 90% of the metagenome for each group. Two-sided Student’s *t*-test with FDR correction. **H)** The abundance differences of the 695 COGs in CN and EU populations based on the ICG database. Two-sided Student’s *t*-test. *, *P*_adj_ < 0.05; **, *P*_adj_ < 0.01; ***, *P*_adj_ < 0.001; n.s., not significant.

We detected universal enrichment of trait COGs across all taxonomic ranks from phylum to genus (Figure 4B), ruling out the possibility that the signal arose from a few large taxa. For the top 20 trait COGs (requiring >20% frequency across all hSGBs, ranked by enriching fold of prevalence), we observed no opposing enrichment trends between different phyla (Figure 4C), supporting that the enrichment resulted from the general selection of host gastrointestinal environment rather than niche differentiation among taxa. Half of the top 20 COGs fall into two functional categories, oxidative stress (*n* = 5) and transporters (*n* = 5).

Because these traits were determined through genomic and metagenomic comparisons, we aimed to investigate whether they represent long-term hSGB evolutionary traits or merely reflect very recent niche adaptation in modern humans (i.e., the rapid change of modern lifestyle). In light of the human “Out of Africa” history, a transitional status in Africans would support these enrichments as more likely long-term evolutionary traits. We thus quantified the relative abundance of the 695 COGs in fecal metagenomes of WNHPs (*n* = 284), non-Africans (*n* = 666), and Africans (*n* = 356, Table S9).^13,22,23, 35–49^ The 356 African metagenomes include samples from the Hadza hunter-gatherers of Tanzania (*n* = 67), who have a remote genetic background to other humans, other rural Tanzanians (*n* = 50), and samples from five other countries (*n* = 239). As shown in Figure 4D, relative abundance and principal coordinate analysis (PCoA) support the transitional status of Africans. Relative abundance of the COGs followed a pattern of non-African human > Hadza ≈ other Tanzanian > WNHP. A similar pattern was observed for other African populations (Fig S4D). These findings suggest that these COGs are more likely co-evolutionary traits due to their successive enriching history in humans.

Given similar COG abundance in Hadza and other rural Tanzanians with distinct lifestyles (hunter-gathering vs. rural), we posited that diet may exert limited effects on the distribution of the trait COGs in the human gut microbiome. Consistent with this hypothesis, we detected no difference in gut metagenomes between vegetarians and omnivores (Figure 4E), nor among two short-term diet-intervention cohorts (Fig S4E&F). Moreover, we also compared the captive primates to wild ones (the same species of *Pan troglodytes* and *Gorilla gorilla*) and found that they have indistinguishable PCoA patterns, with captives even exhibiting a marginally lower abundance of trait COGs than wild counterparts (Figure 4F). However, comparisons of the wild mammalian herbivores, omnivores, and carnivores support the diet-dependent distribution of these COGs in the gut microbiome (Figure 4G). In particular, carnivorous mammals enriched the trait COGs compared with herbivores and omnivores. Moreover, we detected an increasing abundance of these traits in diseased EU and CN individuals (Figure 4H), preliminarily indicating that the trait COGs may negatively correlate with human health, which requires further investigation.

### Enrichment of co-evolutionary traits in gut microbiome is linked with several diseases

We collected 13 datasets examining associations of the gut microbiome with available metagenome and eight diseases (ACVD, atherosclerotic cardiovascular disease, 1 case; NAFLD, nonalcoholic fatty liver disease, 1 case; HTN, hypertension, 1 case; LC, liver cirrhosis, 1 case; CD, Crohn’s disease, 3 cases; OB, obesity, 1 case; RA, rheumatoid arthritis, 1 case; T2D, type 2 diabetes, 2 cases; UC, ulcerative colitis, 2 cases, see Table S10 for detailed information).^39–50^ We selected datasets based on the following criteria: 1) the diseases are strongly related to metabolism or autoimmunity; 2) studies concluding gut microbiome-disease associations; and 3) sound control cohort (in terms of geography, age, *etc*.). Given that 135 of the 695 trait COGs are also enriched in SCEC-hSGBs, which have been implicated in promoting healthy status (Figure 3G), we compared the relative abundance of total and the remaining 560 COGs between the disease and control group for each dataset. Among the 13 datasets, we detected significant differences in four datasets of three diseases (1 ACVD case; 1 NAFLD case; CD, 2 of 3 cases: CD_2 and CD_3), all showing a higher relative abundance of trait COGs in disease versus control groups (*P* < 0.05, two-sided Student’s *t*-test, Figure 5A, see Fig S5A for the other eight datasets). NAFLD only showed an enrichment of 560 COGs, while the others enriched both groups. Figure 5B showed that the COG patterns of disease and control groups diverged in three of the four datasets (in PCoA, permutational multivariate analysis of variance (PERMANOVA), *P* < 0.05). The permutation test excluded that the enrichment of these trait COGs is dependent on overall microbiome divergence between control and disease groups in ACVD, CD_2, and CD_3 (Figure 5C). Furthermore, drug intake may not significantly impact the distribution of the trait COGs (Fig S5B).

**Figure 5.**
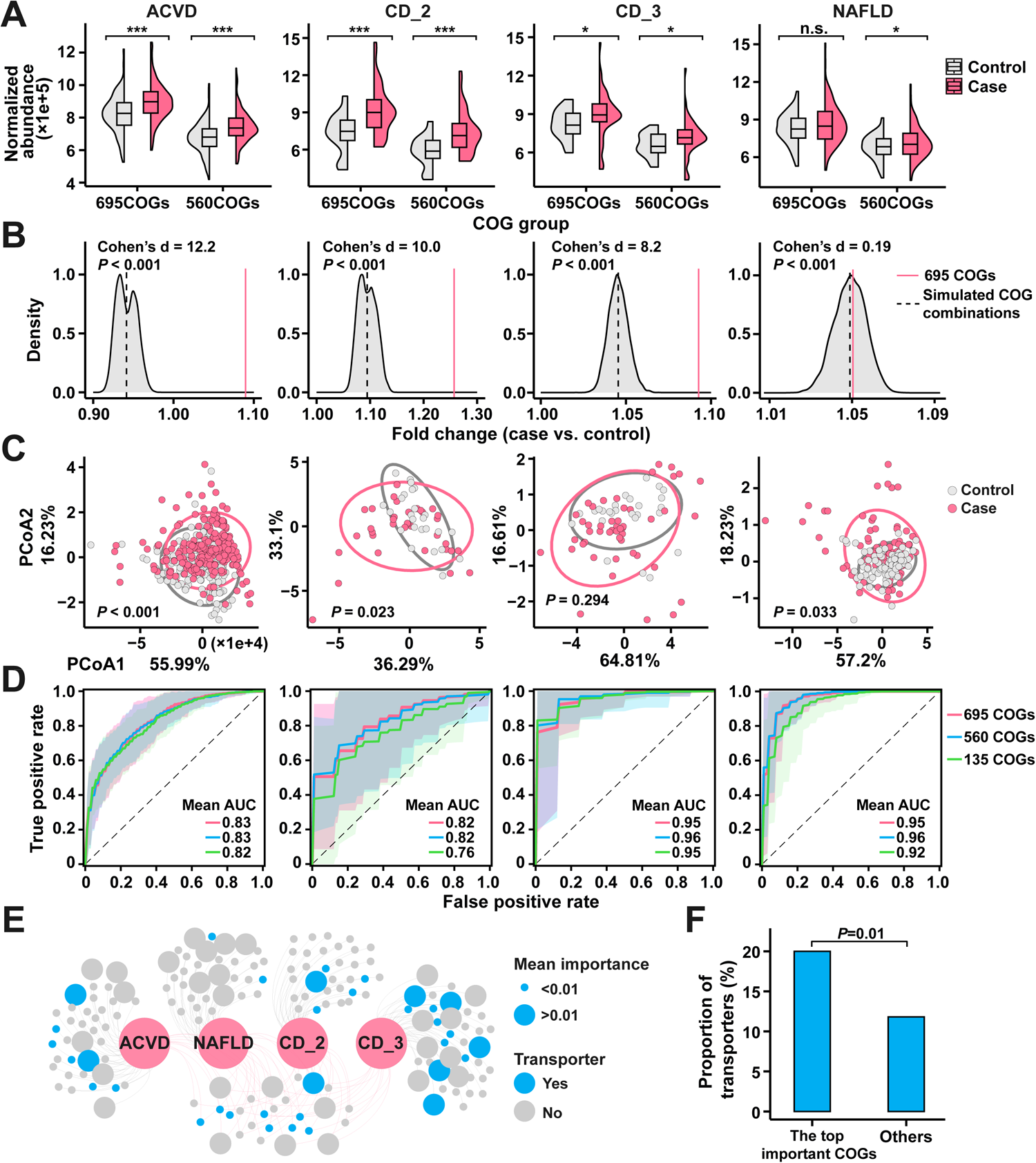
The correlation between 695 evolutionary traits and disease. **A)** The abundance differences of 695 COGs and 560 COGs in the case and control groups. Only four datasets, consisting of three different diseases (ACVD, 1 dataset; NAFLD, 1 dataset; CD, 2 datasets) that exhibited significant enrichment for these traits in the case group, are presented. Two-sided Student’s *t*-test with FDR correction. *P*_adj_ < 0.05; **, *P*_adj_ < 0.01; ***, *P*_adj_ < 0.001; n.s., not significant. **B)** Fold change in case and control groups based on the total abundance of 695 COGs (red solid line) or simulated groups of 695 non-evolutionary traits (black dashed line, *n* = 10,000). One-sample *t*-test. **C)** PCoA based on the Euclidean distance indicated the distribution pattern of 695 COGs in case and control groups. Ellipses cover 90% of the metagenome for each group. PERMANOVA, permutations = 999. **D)** Performance of the random forest classifier based on 695 COGs, 560 COGs, and 135 COGs. The mean AUC and 2-fold standard deviation of 20 bootstraps were shown. **E)** The network depicted the top 50 important COGs in each dataset identified by the random forest classifier based on 695 COGs. The shared COGs are connected by the red line. **F)** The proportion of transporters in the top important traits. Fisher’s exact test.

In addition, our analysis revealed that the trait COGs, regardless of the 695-, 560-, or 135-COGs, had a strong predictive power for host disease in all four datasets, with area under curve (AUC) values ranging from 0.76 to 0.96 (Figure 5D). The 135 COGs exhibited slightly lower AUC values than the other two, possibly due to its limited COG number or their enrichment in SCEC-hSGBs. The top 50 trait COGs with the highest importance during the random forest prediction based on 695 COGs showed no significant overlap among datasets (all *P* >0.05, permutations = 100,000) (Figure 5E). Only 27 traits were shared by more than one dataset, and even for the two CD datasets, the shared top trait COGs were merely 5. These results suggest that the trait COGs are dataset-specific. However, we did observe a higher proportion of transporter COGs distributed in the top trait COGs compared to the others (Figure 5F).

### Defining super enrichers of the co-evolutionary traits and tracking their potential source

The above results suggest that the enrichment of co-evolutionary traits in the gut microbiome is likely linked to several human diseases. We then identified trait COG-enriching hSGBs and investigated their implications for host health. Based on the matrix of the trait COGs, we detected 202 super enricher hSGBs (designated as SUEN-hSGBs) with 2-fold enrichment of the trait COGs (1,203 on average in SUEN vs. 627 in all hSGBs) and three other groups designated as Group A (average encoding COGs: 379), B (average encoding COGs: 601), and C (average encoding COGs: 902) (Figure 6A). Most SUEN-hSGBs are affiliated with Firmicutes, Proteobacteria, and Actinobacteria, but not Bacteroidetes, and exhibit relatively large genome sizes (Figure 6A and Table S11). Remarkably, SUEN-hSGBs are significantly underrepresented among SCEC-hSGBs compared to all hSGBs (16 in 202 vs. 1,342 in 3,779, *P* = 1.17e−20, Fisher’s exact test). In terms of the distribution of the trait COGs, SUEN-hSGBs are not only the generalists with higher coverage (71.2% ± 8.7% vs. 52.9% ± 12.9%) but also the functional enhancers with a higher copy number for detected COGs (2.41 ± 0.48 vs. 1.57 ± 0.24). Among the transporter-related COGs, SUEN-hSGBs were significantly overrepresented compared to Group A and B (Fig S6A). Interestingly, SUEN-hSGBs negatively correlated with Group A and B, which have a low number of trait COGs in their genomes, but slightly positively correlated with Group C containing moderate-enricher (Figure 6B). These results support that the trait COGs are responsible for the niche differentiation of various gut microbial taxa.

**Figure 6.**
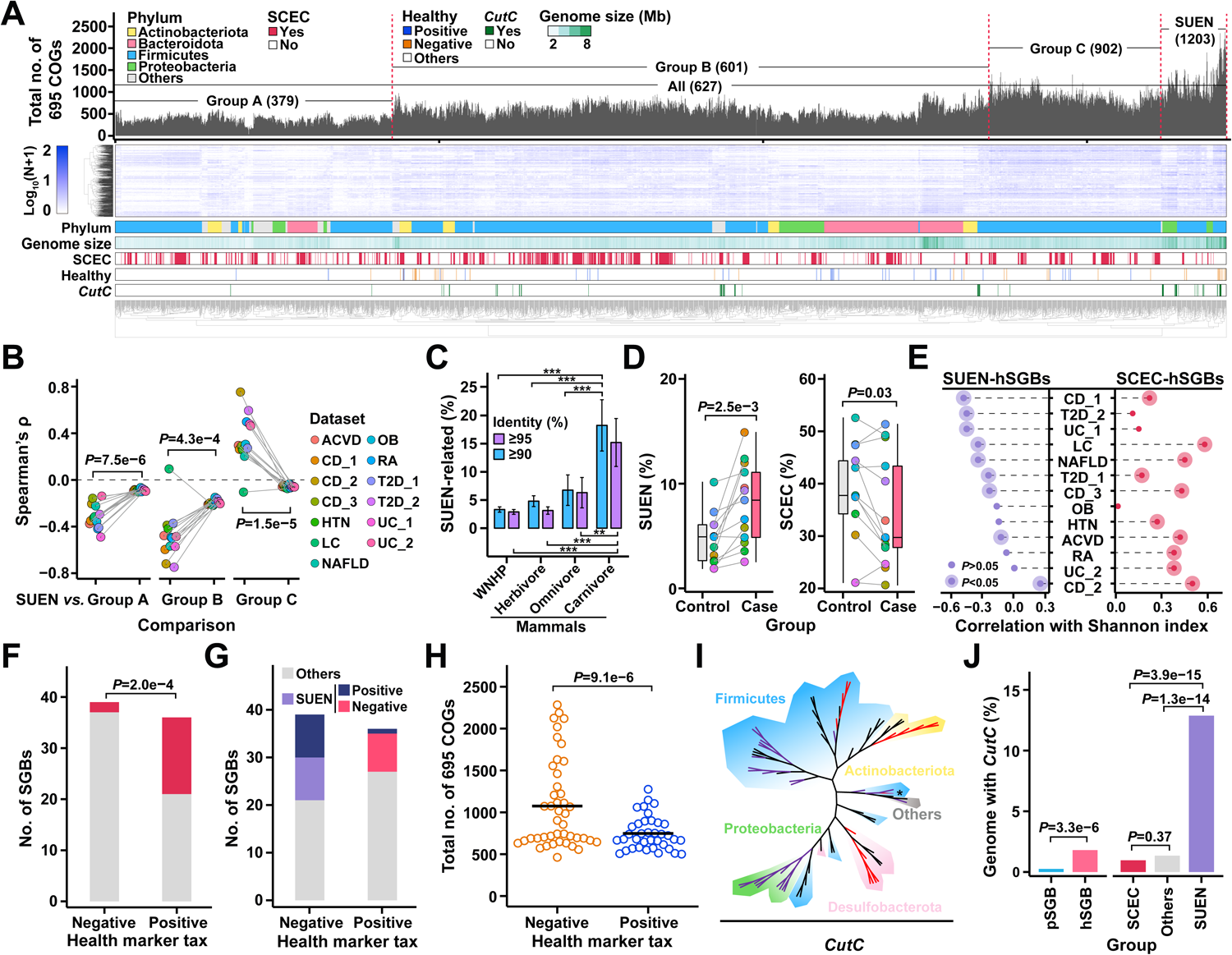
The opposite indications for host health of SUEN-hSGBs and SCEC-hSGBs. **A)** Distribution of 695 co-evolutionary traits in hSGBs (heatmap) and the total copy number of 695 COGs (barplot). The average copy number of 695 COGs in all hSGBs or each group was indicated in parentheses. The COGs (in row) were clustered based the Spearman’s correlation. **B)** Correlation between SUEN-hSGBs and the other three groups in thirteen disease cohorts. Spearman’s ρ between groups in each dataset was calculated based on the proportion of the four groups in each sample. The simulated correlations of each dataset were calculated based on the four counterparts (by group size) that were randomly assigned hSGBs in each sample (*n* = 10,000). Paired two-sided Mann-Whitney U-test. Mean ± s. e. **C)** The proportion of SUEN-related species in WNHP and mammals with different diets. Mean ± s. e. Two-sided Student’s *t*-test with FDR correction. *, *P*_adj_ < 0.05; **, *P*_adj_ < 0.01; ***, *P*_adj_ < 0.001; n.s., not significant. **D)** The proportion of SUEN- and SCEC-hSGBs in control and case groups. Paired two-sided Student’s *t*-test. **E)** Spearman’s ρ between the relative abundance of SUEN- or SCEC-hSGBs with the alpha diversity of the gut microbiome. **F)** A higher proportion of SCEC-hSGBs were detected in healthy-positive species. Fisher’s exact test. **G)** The distribution of SUEN-hSGBs and their related species in healthy indicated species. **H)** The total copy number of 695 COGs in healthy indicating species. Two-sided Student’s *t*-test. **I)** Maximum-likelihood phylogenetic tree of *CutC*. The SGBs affiliated with SCEC or SUEN were labeled by branch colors (red, SCEC; purple, SUEN; *, both). **J)** The proportion of SGBs encoding *CutC* in the two databases and SCEC- or SUEN-hSGBs. Fisher’s exact test.

Given the low proportion of the SUEN-hSGBs affiliated with SCECs, we are interested in their source. We hypothesize that SUEN-hSGBs partially transferred from other mammals, as an enrichment of the trait COGs in carnivorous mammals was observed (Figure 4G). To test this, we profiled the distribution of SUEN-hSGBs-related taxa in the gut metagenomes of WNHPs, wild herbivorous, omnivorous, and carnivorous mammals using a taxonomic marker gene (ribosomal protein L1, COG0081) (see Methods). The SUEN-hSGB-related taxa (≥ 90% or 95% amino acid identity for metagenomic reads) were much more abundant in carnivores than in WNHPs, herbivores and omnivores (All *P*<0.01, two-sided Student’s *t*-test, Figure 6C). Thus, we propose that carnivorous mammals were potential sources of some SUEN-hSGBs, although the detailed history of transfer and diversification remains unclear.

### SUEN- and SECE-hSGBs have opposite implications for gut microbiome dysbiosis and human health

We then investigated the potential implications of the SUEN-hSGBs and SCEC-hSGBs, which have shown a decreasing trend in diseased individuals (Figure 3G), for host health. Firstly, we determined their relative abundances in the 13 datasets. As shown in Figure 6D, SUEN-hSGBs and SCEC-hSGBs were significantly enriched in diseased and healthy individuals, respectively, although their relative abundances varied greatly among datasets.

Secondly, we determined whether there was a correlation between these taxa and the alpha diversity of the gut microbiome, which is a common indicator of dysbiosis.^51,52^ As shown in Figure 6E, the Shannon index for most 13 datasets positively correlated with SCEC-hSGBs (10 of 13, *P* < 0.05) but negatively with SUEN-hSGBs (8 of 13, *P* < 0.05). Even those non-significant correlations were consistent in their direction with only one exception (SUEN in CD_2, Spearman’s ρ = 0.26). The Shannon indexes were calculated by removing either SCEC-hSGBs or SUEN-hSGBs. Alternative statistics including these taxa yielded similar results (Fig S6B). The above results indicated that the abundance of SCEC-hSGBs and SUEN-hSGBs oppositely correlate with the diversity of the gut microbiome.

Thirdly, since two previous studies have provided the list of hSGBs positively or negatively related to general human health based on large cohorts, we then investigated how the SCEC-hSGBs and SUEN-hSGBs are involved in these marker taxa (36 for health-positive and 39 for health-negative, see Table S12 for the taxonomic information).^53,54^ The results showed that SCEC-hSGBs were more biased towards health-positive hSGBs (*P* < 0.001, Fisher’s exact test, Figure 6F), whereas SUEN-hSGBs showed the opposite pattern (Figure 6G). Moreover, we found that other hSGBs positively correlating with the total SUEN-hSGBs in the metagenomes were also significantly related to health-negative hSGBs (8 and 1 for health-negative and health-positive hSGBs, respectively), while negatively correlating hSGBs were only health-positive (*n* = 9). In addition, the sum of the trait COGs was significantly higher in health-negative hSGBs than in health-positive ones (Figure 6H).

Lastly, given the strong correlation or even causation between gut microbial metabolites trimethylamine (TMA) and trimethylamine-N-oxide (TMAO) and several diseases such as ACVD, NAFLD, and IBD,^40,55,56^ we analyzed *CutC* (the choline-TMA-lyase) in hSGBs, since it is the primary enzyme responsible for generating TMA.^57^ Consistent with the previous report,^58^ we found that the prevalence of *CutC*-encoding hSGBs is rare (68 in 3,779, Figure 6I&J), with even lower pSGB prevalence (Figure 6J). We confirmed almost no intraspecific variation in encoding *CutC* (Fig S6C). In the phylogenetic tree of *CutC* from hSGBs, the proteobacterial *CutC*, which form a unique clade and all belong to Enterobacteriaceae, were exclusively found in SUEN-hSGBs (Figure 6I). Notably, the relative abundance of *CutC*-encoding Enterobacteriaceae in fecal metagenome is associated with urinary TMAO level,^59^ suggesting the related taxa are responsible for TMA production. Moreover, the frequency of encoding *CutC* is approximately 10-fold higher in SUEN-hSGBs than in the other hSGBs, whereas no significant difference was observed for the SCEC-hSGBs with the background (Figure 6I).

Together, these results indicated SCEC-hSGBs and SUEN-hSGBs had opposite implications on host health.

## Discussion

In this study, we established a genomic database of gut microbial species from WNHPs to define co-evolutionary species and traits of hSGBs. It is important to note that the pSGBs database is obviously unsaturated because there are over 500 extant primate species,^60^ which may have intraspecific divergence in their gut microbiome.^61^ Therefore, a more comprehensive collection of samples from primates will improve the reliability of the list of co-evolving species and evolutionary traits. However, the rarefaction curve shows that increasing SCEC-hSGBs are approaching saturation (Fig S1B), indicating the current pSGB database fairly represents co-evolutionary lineages.

Few pSGBs are shared across wild primates and human, suggesting geographical isolation dominates wild primate gut microbiome histories, at least for a short term. This confirms the irreplaceability of wild animal gut microbiome studies^14^ and potentially supports allopatric speciation as a major driving force of gut microbe-host co-speciation.^4^ However, many hSGBs show host-jumping events, which may occur in the long-run evolutionary history. The *Homo* lineage has increased carnivory over 2 million years relative to other primates,^62^ which may increase the chance of transferring gut microbes from the primate preys, leading to the initiation of co-evolution with the new host.

The evolutionary trend for hSGBs can be observed at different timescales. For a very short period, such as within a host lifespan or even a few years, evidence has shown detectable mutation and gene gain/loss events that suggest adaptations.^63,64^ As the timescale increases slightly, strong selective functional potentials can introduce apparent adaptive changes in the genome.^13^ For example, our results show an increased prevalence of ARGs in hSGBs following <100 years of the corresponding antibiotic usage. Other studies have demonstrated that population-level dietary significantly impacts the intraspecific CAZy profiles of hSGBs.^65^ On the long-term co-evolutionary scale across host species, our study observed an evolutionary trend of genomic reduction for SCEC-hSGBs. Extreme genome reduction is well-known in symbiotic bacteria compared with their free-living relatives.^66,67^ However, the genomic reduction in SCEC-hSGBs, although to a lesser extent, was based on a comparison with corresponding SCEC-pSGBs, which are also symbionts. This can be an adaptive outcome of the putative higher stability of the human gut nutritional condition compared to that of wild primates. Supporting this, lost genes were biased towards cell motility and energy production functions. Decreasing chemistry complexity of food, reflected in the simplified GH families in hSGBs, as well as the putatively increased host digestive capability (much lower stomach pH in human than primates),^68^ may also drive the reduction.

Although the enriching traits of hSGBs compared to pSGB were functionally diverse, a hallmark of hSGBs is those adaptive to oxidative stress, suggesting higher intestinal oxidative toxicity in humans than in wild primates. Lifestyle factors like high-fat diets and sleep deprivation increase gut ROS.^69,70^ While human diets contain more fat than wild primates,^71^ comparisons of vegetarian and omnivorous humans and results of short-term high-fat diet studies do not support diet-induced enrichment of trait COGs. In contrast, we found the traits enriched in carnivorous vs. omnivorous or herbivorous mammals. One possible explanation for this contradiction is that only long-term dietary changes or extreme short-term changes, and permanent dietary shifts that induced irreversible host physiological alterations are responsible for the enrichment of trait COGs. As it has been known that the trophic level can extensively link with adaptive changes in host physiological and metabolic features.^68,72^

The co-evolutionary lineages in hSGBs positively correlated with host health, consistent with the hypothesis that long-term co-evolution tends to select mutualisms instead of antagonisms.^73^ This is also consistent with the proposal that the loss of specific bacterial species from our ancestral microbiome could result in an increased risk of chronic diseases.^74^ A previous study based on the rRNA genes showed that human gut bacterial genera containing more co-speciating taxa across mammals were negatively correlated with IBD,^7^ suggesting links between the long-term co-evolving bacteria and host immunity. Our results showed that the SCEC-hSGBs strongly positively correlated with the alpha diversity of the gut microbiome, indicating an association with eubiosis, though the causality is unclear.

In contrast, the enrichment of the co-evolutionary traits in the gut microbiome is associated with certain human diseases. The super enrichers of these traits positively correlated with host unhealth and dysbiosis. Moreover, besides correlation, disproportionate TMA producers associated with SUEN-hSGBs have the potential to cause specific diseases as the metabolites have a significant impact on human health.^75^ Notably, the SUEN-hSGBs contain a biasedly lower proportion of SCEC ones, indicating most of them are outsiders that transferred into the human gut more recently. Our preliminary analysis suggested carnivorous mammals as a potential source of some SUEN-hSGBs, though transfer histories remain unprofiled. The much higher *CutC* frequency in SUEN-hSGBs further supports the hypothesis, as choline, the substrate of *Cuts*, is more abundant in animal tissues than in plants. More importantly, the disproportional and key TMA producers affiliated with SUEN-hSGBs suggest their potential role as causative agents for certain diseases, considering the direct impact of the metabolites on human health.^75,76^

We posit that the prevalence of these super enrichers can be at least partially explained by niche selection. It is supported by the fact that their co-occurring taxa are more likely traits enrichers and negatively correlated taxa encoding fewer trait COGs, strongly suggesting the trait COGs play certain roles in their niche adaptation. Within the traits, COGs of transporters are highly represented in SUEN-hSGBs. It has been reported that encoding redundant transporters can increase the fitness of gut bacteria.^77^ According to our results, the enrichment of transporters also seemed related to several diseases according to the random forest predictive results. Moreover, a previous study has proposed generalists with larger genomes have advantages in unstable environments.^78^ The SUEN-hSGBs with larger genome sizes may convey heightened competitive ability against other taxa, including SCEC-hSGBs, in both pre-disease and disease conditions. The loss of SCEC-hSGBs and over-enrichment of the traits in the gut microbiome reflect (or are selected by) the host gastrointestinal status that deviates from evolutionary and ecological normality.

In conclusion, our study characterized long-term co-evolutionary lineages and traits of human gut microbe compared to an updated gut microbial genome database of WNHP and revealed their opposing correlations with the host’s health status. The SUEN- and SCEC-hSGBs may serve as new biomarkers, beyond those obtained from cohort studies, for predicting and diagnosing host health or disease. Defining the SCEC-hSGBs may provide valuable guidance for developing probiotics and other potential gut microbial resources, as they are theoretically safer and better adapted to the host.

### Star methods

- Public data collection
- Fecal sample collection and metagenomic sequencing
- Genome reconstruction and species-level genome clustering
- Taxonomy assignment and phylogenetic analysis
- Functional annotation
- SCEC definition
- Mapping the hSGBs to IGC
- Co-evolutionary traits definition
- Quantification functional genes in metagenomes
- Random forest classifier
- Network analysis
- Host’s phylogenetic group, diet, and divergence time of NHPs.
- Statistical analysis and data visualization
- Data and code availability

## Acknowledgments

This work received finical support from National Natural Science Foundation of China (32270006). We thank Wuyishan National Nature Reserve, the Zoo of Xiamen and the Zoo of Nanping for fecal sample collection of NHPs.

## Author contributions

Conceptualization, H. L. and F. G.; methodology, H. L. and F. G.; investigation, H. L. and J. H.; resources, H. L., J. H., J. L., W. Z., and Y. L.; writing – original draft, H. L. and F. G.; writing – review & editing, H. L., Q. Q., and F. G.; funding acquisition, F. G.

## Materials and methods

### Public data collection

We collected 321 public metagenomes spanning 23 WNHPs for an updated genomic database of NHP.^17,23–30^ In addition, we also collected 2,096 fecal metagenomes of humans, including 1,890 from studies on gut microbiome-metabolism/autoimmunity correlations,^39–50^ 356 from Africans,^13,22,23, 35–38^ and 116 from diet investigation or intervention.^65,79,80^ For the disease cohorts, we only selected one metagenome per individual as a representative. Furthermore, we also collected 91 metagenomes from 71 mammals with different diets.^81^ Details are in Tables S1, S9, and S10.

We referred to publicly available prokaryotic genomes from three studies, including 1) the human gut MAG/SGB database constructed by Pasolli et al.(2019),^22^ with MAGs and SGBs from other body sites removed; 2) the genomic database of NHPs;^16^ 3) the reference genomes from GTDB database r95.^31^

In addition, we referred to 81 gut bacterial species associated with human health or disease determined in two large cohort studies.^53,54^ Given that species names may be inconsistent across taxonomic systems, for these species, we selected a representative genome from NCBI and then determined its corresponding representative genomes in hSGB using fastANI (v1.3, ANI >95%; see Table S12 for details).^33^

### Fecal sample collection and metagenomic sequencing

We collected fecal samples from wild *M. thibetana* (*n* = 12) in Wuyishan National Nature Reserve and Da’anyuan, Nanping, Fujian province, China. Additionally, we collected fecal samples of captive *M. mulatta* (*n* = 6), *M. mulatta* (*n* = 4), and *M. thibetana* (*n* = 3) from the Zoo of Xiamen and Nanping, Fujian province, China. The fresh feces were collected in sterile collection tubes containing 70% ethanol. All samples were shipped with dry ice and stored at −80℃ freezer until use. DNA of 25 samples collected in this study was extracted using FastDNA® SPIN Kit for Feces (MP, USA) DNA extraction kit. The concentration and quality of DNA were determined by NanoDrop and agarose gel electrophoresis, respectively. The metagenomic library was constructed using NEBNext® Ultra DNA Library Prep Kit for Illumina (NEB, USA). The samples were sequenced with the PE150 strategy on Illumina Hiseq Novaseq 6000 platform (commercial service, Novogene, Beijing).

### Genome reconstruction and species-level genome clustering

We filtered the low-quality reads from all metagenomes of NHPs using Trimmomatic v.0.38^82^ and assembled the filtered reads using metaSPAdes (v.3.9.1, parameters: -k 33, 45, 55; for paired-end sample) or MEGAHIT (v1.1.4, for single-end or metagenomes with bases >15 Gb).^83,84^ We binned scaffolds >1,000bp using MetaWRAP v1.2.1 with two binners (MaxBin2 and metaBAT2, default parameters),^85–87^ and refined MAGs using the bin_refinement module. CheckM (v1.0.7; lineage-specific workflow) was used to estimate the quality of MAGs and only those with genome completeness >50% and contamination <5% were kept.^88^ MAGs were clustered using dRep (v2.6, parameter: -p 50 -ignore genomequality -pa 0.70 -sa 0.95 --S_algorithm fastANI) at the threshold of 95% ANI by two-step cluster.^89^ The 2,036 MAGs with the best quality of each SGB cluster were chosen as the representative genomes. We dereplicated the MAGs from NHP2019 using the same pipeline, resulting in 1,232 SGB clusters.

We aligned the filtered reads of the gut metagenomes from WNHPs to the contigs of 4,942 MAGs using Bowtie2 v2.3.4.3 with default parameters,^90^ and calculated the mapping rate by dividing the total mapped reads by all quality-filtered reads of each sample.

### Taxonomy assignment and phylogenetic analysis

Taxonomy assignment for MAGs and SGBs was determined by GTDB-Tk (v1.3.0; ‘classify_wf’ workflow and default parameters) based on the GTDB database (release 95).^31,91^ Phylogenetic analyses of 1,637 bacterial pSGBs based on concatenation of 120 ubiquitous single-copy genes.^31^ The 120 markers were extracted from the annotation results of GTDB-Tk and were aligned using Mafft v7.407.^92^ Genomes with >60% gaps in the concatenated alignment were removed. The phylogenetic tree was inferred using FastTree v2.1 under the WAG + GAMMA models and visualized via the iTOL.^93,94^

### Functional annotation

The open reading frames (ORFs) of pSGBs and hSGBs were predicted using Prodigal v2.6.3.^95^ The functional profile of each SGB was performed using eggNOG-mapper (v2.1.6, -m diamond) with eggnog database v5.0 under default parameters.^96,97^ CAZymes were annotated using the run_dbcan,^98^ and the substrates categories of the top 5 CAZy families enriched in either pSGB or hSGB were manually retrieved from the *nr* database.^99^ ARGs were annotated using DeepARG v2.^100^

For *CutC* (encoding the choline trimethylamine-lyase) annotation, a total of 1,184 proteins affiliated with K20038 (KEGG ortholog, choline trimethylamine-lyase) from pSGBs and hSGBs were annotated against the *nr* database using BLASTP (evalue <1e−5, -max_target_seqs 100).^99^ Alignments with annotation targeted to choline trimethylamine-lyase were filtered with identity >45% and coverage >50%.^58^ Filtered sequences were aligned using Mafft v7.407,^92^ and the phylogenetic tree was inferred using RAxML v8.2.12 with the parameters ‘-# 100 -m PROTGAMMAAUTO --auto-prot=aic’.^101^ Finally, 73 protein sequences from 72 SGBs that formed a monophyletic branch were determined as *CutC*. To evaluate the intraspecific divergence of *CutC*, up to 100 high-quality MAGs were randomly selected for 56 hSGB clusters (56 *CutC* encoding species). Protein sequences were aligned against the 73 *CutC* sequences using BLASTP (evalue <1e−5),^99^ and alignments were filtered with identity >90% and coverage >90%.

### SCEC definition

In this study, we used an ANI-based method to define co-evolutionary clusters. To determine the operational threshold for co-evolutionary clusters, we calculated the split ratio for non-singleton SGB clusters by stepwise increasing the ANI cutoff (from 70% to 95%, 1% per step) using dRep (v2.6.2, cluster module; options ‘--clusterAlg average --S_algorithm fastANI --cov_thresh 0.1’).^89^ Only 107 genera with ≥10 SGBs were considered to guarantee enough non-singleton clusters. For instance, there are 309 non-singleton clusters under the ANI cutoff at 70%, and 8 of them split after using the cutoff at 71%. Therefore, the split ratio for ANI-71% is 2.6%. Fisher’s exact test was for identifying the first significant increase in split ratio. Finally, after combining all pSGBs and hSGBs, the co-evolutionary clusters were determined using the cutoff ANI-77%. Therein, those containing both pSGB and hSGBs were referred to as SCEC ones.

### Co-evolutionary traits definition

To identify co-evolutionary traits that enriched in hSGB, the Mann-Whitney U-tests (abundance-based) and Fisher’s exact test (prevalence-based) were used for comparing 1,635 high-frequency (frequency >20%) COGs between interspecific hSGBs and pSGBs. We determined 839 COGs significantly enriched in hSGB (hSGB versus pSGB >1 and FDR-corrected *P* <0.05) using both methods. Among them, 695 COGs as co-evolutionary traits because they showed significant enrichment in metagenomes of healthy populations compared to WNHP.

### Quantification of functional genes in metagenomes

Considering the under-representation of pSGBs in the widely referred genome databases, we developed a pipeline to quantify the relative abundance of functional genes and SGB in metagenomes (Fig S4A, B, and C). Firstly, we de-replicated annotated ORFs from pSGBs and hSGBs with 95% identity and 90% coverage using CD-HIT v4.7,^102^ including representatives in our customized database. Secondly, we subsampled quality-filtered metagenomes to 10 million reads to reduce the computational burden and minimize any potential distortion caused by unequal sequencing depth. We included all reads for metagenomes <10 million reads and removed metagenomes <1 million reads. Thirdly, the reads were aligned against the customized database using DIAMOND BLASTX (-evalue <1e−5, -max_target_seqs = 1),^103^ and alignments are filtered with identity >50% and coverage >80%. The filtered result was normalized to per million reads for each metagenome. Finally, we observed high annotation rate variation between metagenomes, potentially from non-prokaryotic DNA proportions, so we normalized using the ribosomal protein L1.

Our pipeline achieved higher annotation rates for WNHP and human gut metagenomes than the HUMAnN3 (uniref90_201901b_full database, default parameters, Fig S4B).^104^ Correlation coefficients among three conserved proteins (ribosomal protein L1, L3 (COG0087), and S3 (COG0092)) were also higher in our pipeline than in the HUMAnN3 results (Fig S4C).

To quantify the SUEN-related taxa in metagenomes of WNHPs, wild herbivorous, omnivorous, and carnivorous mammals, we aligned the reads to the COG0081 database from hSGB and pSGBs using DIAMOND BLASTX. The alignments were filtered with ≥90% or 95% amino acid identity and >80% coverage and then normalized to per million reads for each metagenome. For each threshold, the abundance of SUEN-related taxa is characterized by the ratio of the number of reads mapped to SUEN-hSGBs to the total number of reads mapped to all SGBs under identity ≥50%.

For quantifying hSGBs in metagenomes, the relative abundance of each species is computed by totaling the relative abundance of 120 universal single-copy genes.

### IGC data analysis

To define the proportion of SCEC-hSGBs in different regions, the ribosomal protein genes (ribosomal protein L1) in the IGC database were aligned against the corresponding ones of the hSGBs using BLASTN,^34,99^ and alignments were filtered with identity >95% and coverage >90%. The filtered sequences were labeled as SCEC-hSGB, other-hSGB (based on the respective hSGB group), or others (sequences without hits). The proportion of the CN and EU populations was calculated for each metagenome based on the gene-level relative abundance table offered by ICG.

### Random forest classifiers

We built random forest classifiers using scikit-learn to evaluate trait COG performance in predicting case/control groups.^105^ The dataset was randomly split into training and test sets (7:3) 20 times for each cohort, and the model was trained using optimized parameters to achieve the predicted performance. Mean AUC was used to evaluate the performance of the classifier. The top 50 COGs, determined by importance, were subsequently analyzed.

### Network analysis

Correlations of hSGBs were calculated using FastSpar v1.0.0 based on the relative abundance of a combination of 13 disease datasets.^106^ Notably, due to the unbalanced sample size of these studies, the metagenome of ACVD and T2D_1 were randomly sampled to 200 (100 samples each in the control and case group) to reduce computational bias. To reduce computational effort, species with a prevalence <10% were excluded from the analysis. We used permutation testing (*n* = 5,000) and Benjamini-Hochberg correction for multiple testing to generate *P* values. The network was visualized using Cytosacpe v3.9.1.^107^

### Phylogenetic group, diet, and divergence time of NHPs

The phylogeny and divergence time of primates are retrieved from Timetree (http://timetree.org/). The dietary of primates is collected from Animal Diversity Web (https://animaldiversity.org/).

### Statistical analysis and data visualization

We calculated alpha diversity using the Shannon index based on hSGB relative abundance in metagenome (Vegan R package).^108^ The PCoA was performed using the vegan R package based on the normalized relative abundance matrix of 695 co-evolutionary traits in each metagenome. The difference in clustering patterns based on 695 co-evolutionary traits between the control and case groups was tested using permutational analysis of variance. Significances of the shared COGs of the top important traits between the four datasets were inferred using simulated sampling based on the multivariate hypergeometric distribution (permutations = 100,000; Purrr R package).

We performed statistics in R v4.1.3. We report Student’s *t*-tests, Mann-Whitney U tests, Fisher’s exact tests, Chi-Square tests, and Benjamini-Hochberg false discovery rate corrections for multiple hypothesis testing.

### Data and code availability

The raw sequencing data of non-human primates in this study are available in the Sequence Read Archive (SRA) under Bioproject PRJNA932532. MAGs recovered in this study are available in the FigShare repository (https://doi.org/10.6084/m9.figshare.22117169).

## Reference

1. Sommer, F., and Bäckhed, F. (2013). The gut microbiota—masters of host development and physiology. Nat. Rev. Microbiol. 11, 227–238.

2. Tremaroli, V., and Bäckhed, F. (2012). Functional interactions between the gut microbiota and host metabolism. Nature 489, 242–249.

3. Davenport, E.R., Sanders, J.G., Song, S.J., Amato, K.R., Clark, A.G., and Knight, R. (2017). The human microbiome in evolution. BMC Biol. 15, 1–12.

4. Groussin, M., Mazel, F., and Alm, E.J. (2020). Co-evolution and co-speciation of host-gut bacteria systems. Cell Host Microbe 28, 12–22.

5. Janzen, D.H. (1980). When is it coevolution? Evolution 34, 611–612.

6. Moran, N.A., and Sloan, D.B. (2015). The hologenome concept: helpful or hollow? PLoS Biol. 13, e1002311.

7. Groussin, M., Mazel, F., Sanders, J.G., Smillie, C.S., Lavergne, S., Thuiller, W., and Alm, E.J. (2017). Unraveling the processes shaping mammalian gut microbiomes over evolutionary time. Nat. Commun. 8, 14319.

8. Rojas, C.A., Ramírez-Barahona, S., Holekamp, K.E., and Theis, K.R. (2021). Host phylogeny and host ecology structure the mammalian gut microbiota at different taxonomic scales. Animal Microbiome 3, 1–18.

9. Lim, S.J., and Bordenstein, S.R. (2020). An introduction to phylosymbiosis. Proc. R. Soc. B 287, 20192900.

10. Mallott, E.K., and Amato, K.R. (2020). Phylosymbiosis, diet and gut microbiome-associated metabolic disease. Evol. Med. Public Hlth. 2020, 100–101.

11. Li, H., Meier-Kolthoff, J.P., Hu, C., Wang, Z., Zhu, J., Zheng, W., Tian, Y., and Guo, F. (2022). Panoramic insights into microevolution and macroevolution of a *Prevotella copri*-containing lineage in primate guts. Genomics Proteomics Bioinformatics 20, 334–349.

12. Moeller, A.H., Caro-Quintero, A., Mjungu, D., Georgiev, A.V., Lonsdorf, E.V., Muller, M.N., Pusey, A.E., Peeters, M., Hahn, B.H., and Ochman, H. (2016). Cospeciation of gut microbiota with hominids. Science 353, 380–382.

13. Suzuki, T.A., Fitzstevens, J.L., Schmidt, V.T., Enav, H., Huus, K.E., Mbong Ngwese, M., Grießhammer, A., Pfleiderer, A., Adegbite, B.R., Zinsou, J.F., et al. (2022). Codiversification of gut microbiota with humans. Science 377, 1328–1332.

14. Amato, K.R., Yeoman, C.J., Kent, A., Righini, N., Carbonero, F., Estrada, A., Rex Gaskins, H., Stumpf, R.M., Yildirim, S., Torralba, M., et al. (2013). Habitat degradation impacts black howler monkey (*Alouatta pigra*) gastrointestinal microbiomes. ISME J. 7, 1344–1353.

15. Clayton, J.B., Vangay, P., Huang, H., Ward, T., Hillmann, B.M., Al-Ghalith, G.A., Travis, D.A., Long, H.T., Tuan, B.V., Minh, V.V., et al. (2016). Captivity humanizes the primate microbiome. Proc. Natl. Acad. Sci. USA 113, 10376–10381.

16. Manara, S., Asnicar, F., Beghini, F., Bazzani, D., Cumbo, F., Zolfo, M., Nigro, E., Karcher, N., Manghi, P., Metzger, M.I., et al. (2019). Microbial genomes from non-human primate gut metagenomes expand the primate-associated bacterial tree of life with over 1000 novel species. Genome Biol. 20, 299.

17. Amato, K.R., G. Sanders, J., Song, S.J., Nute, M., Metcalf, J.L., Thompson, L.R., Morton, J.T., Amir, A., J. McKenzie, V., Humphrey, G., et al. (2019). Evolutionary trends in host physiology outweigh dietary niche in structuring primate gut microbiomes. ISME J. 13, 576–587.

18. Walter, J., and Ley, R. (2011). The human gut microbiome: ecology and recent evolutionary changes. Annu. Rev. Microbiol. 65, 411–429.

19. Corbett, S., Courtiol, A., Lummaa, V., Moorad, J., and Stearns, S. (2018). The transition to modernity and chronic disease: mismatch and natural selection. Nat. Rev. Genet. 19, 419–430.

20. Eaton, S.B., Konner, M., and Shostak, M. (1988). Stone agers in the fast lane: chronic degenerative diseases in evolutionary perspective. Am. J. Med. 84, 739–749.

21. Sonnenburg, E.D., and Sonnenburg, J.L. (2019). The ancestral and industrialized gut microbiota and implications for human health. Nat. Rev. Microbiol. 17, 383–390.

22. Pasolli, E., Asnicar, F., Manara, S., Zolfo, M., Karcher, N., Armanini, F., Beghini, F., Manghi, P., Tett, A., Ghensi, P., et al. (2019). Extensive unexplored human microbiome diversity revealed by over 150,000 genomes from metagenomes spanning age, geography, and lifestyle. Cell 176, 649–662. e20.

23. Campbell, T.P., Sun, X., Patel, V.H., Sanz, C., Morgan, D., and Dantas, G. (2020). The microbiome and resistome of chimpanzees, gorillas, and humans across host lifestyle and geography. ISME J. 14, 1584–1599.

24. D’arc, M., Ayouba, A., Esteban, A., Learn, G.H., Boué, V., Liegeois, F., Etienne, L., Tagg, N., Leendertz, F.H., Boesch, C., et al. (2015). Origin of the HIV-1 group O epidemic in western lowland gorillas. Proc. Natl. Acad. Sci. USA 112, E1343–E1352.

25. Greene, L.K., Williams, C.V., Junge, R.E., Mahefarisoa, K.L., Rajaonarivelo, T., Rakotondrainibe, H., O’Connell, T.M., and Drea, C.M. (2020). A role for gut microbiota in host niche differentiation. ISME J. 14, 1675–1687.

26. Hicks, A.L., Lee, K.J., Couto-Rodriguez, M., Patel, J., Sinha, R., Guo, C., Olson, S.H., Seimon, A., Seimon, T.A., Ondzie, A.U., et al. (2018). Gut microbiomes of wild great apes fluctuate seasonally in response to diet. Nat. Commun. 9, 1786.

27. Orkin, J.D., Webb, S.E., and Melin, A.D. (2019). Small to modest impact of social group on the gut microbiome of wild Costa Rican capuchins in a seasonal forest. Am. J. Primatol. 81, e22985.

28. Tsukayama, P., Boolchandani, M., Patel, S., Pehrsson, E.C., Gibson, M.K., Chiou, K.L., Jolly, C.J., Rogers, J., Phillips-Conroy, J.E., and Dantas, G. (2018). Characterization of wild and captive baboon gut microbiota and their antibiotic resistomes. mSystems 3, e00016–00018.

29. Tung, J., Barreiro, L.B., Burns, M.B., Grenier, J.-C., Lynch, J., Grieneisen, L.E., Altmann, J., Alberts, S.C., Blekhman, R., and Archie, E.A. (2015). Social networks predict gut microbiome composition in wild baboons. eLife 4, e05224.

30. Shaffer, J.P., Nothias, L.-F., Thompson, L.R., Sanders, J.G., Salido, R.A., Couvillion, S.P., Brejnrod, A.D., Lejzerowicz, F., Haiminen, N., Huang, S., et al. (2022). Standardized multi-omics of Earth’s microbiomes reveals microbial and metabolite diversity. Nat. Microbiol., 1-23.

31. Parks, D.H., Chuvochina, M., Chaumeil, P.-A., Rinke, C., Mussig, A.J., and Hugenholtz, P. (2020). A complete domain-to-species taxonomy for Bacteria and Archaea. Nat. Biotechnol. 38, 1079–1086.

32. Gosselin, S., Fullmer, M.S., Feng, Y., and Gogarten, J.P. (2022). Improving phylogenies based on average nucleotide identity, incorporating saturation correction and nonparametric bootstrap support. Syst. Biol. 71, 396–409.

33. Jain, C., Rodriguez-R, L.M., Phillippy, A.M., Konstantinidis, K.T., and Aluru, S. (2018). High throughput ANI analysis of 90K prokaryotic genomes reveals clear species boundaries. Nat. Commun. 9, 5114.

34. Li, J., Jia, H., Cai, X., Zhong, H., Feng, Q., Sunagawa, S., Arumugam, M., Kultima, J.R., Prifti, E., Nielsen, T., et al. (2014). An integrated catalog of reference genes in the human gut microbiome. Nat. Biotechnol. 32, 834–841.

35. Rampelli, S., Schnorr, S.L., Consolandi, C., Turroni, S., Severgnini, M., Peano, C., Brigidi, P., Crittenden, A.N., Henry, A.G., and Candela, M. (2015). Metagenome sequencing of the Hadza hunter-gatherer gut microbiota. Curr. Biol. 25, 1682–1693.

36. Smits, S.A., Leach, J., Sonnenburg, E.D., Gonzalez, C.G., Lichtman, J.S., Reid, G., Knight, R., Manjurano, A., Changalucha, J., Elias, J.E., et al. (2017). Seasonal cycling in the gut microbiome of the Hadza hunter-gatherers of Tanzania. Science 357, 802–806.

37. Tett, A., Huang, K.D., Asnicar, F., Fehlner-Peach, H., Pasolli, E., Karcher, N., Armanini, F., Manghi, P., Bonham, K., Zolfo, M., et al. (2019). The *Prevotella copri* complex comprises four distinct clades underrepresented in westernized populations. Cell Host Microbe 26, 666–679. e7.

38. Rubel, M.A., Abbas, A., Taylor, L.J., Connell, A., Tanes, C., Bittinger, K., Ndze, V.N., Fonsah, J.Y., Ngwang, E., Essiane, A., et al. (2020). Lifestyle and the presence of helminths is associated with gut microbiome composition in Cameroonians. Genome Biol. 21, 122.

39. Franzosa, E.A., Sirota-Madi, A., Avila-Pacheco, J., Fornelos, N., Haiser, H.J., Reinker, S., Vatanen, T., Hall, A.B., Mallick, H., McIver, L.J., et al. (2019). Gut microbiome structure and metabolic activity in inflammatory bowel disease. Nat. Microbiol. 4, 293–305.

40. Jie, Z., Xia, H., Zhong, S.-L., Feng, Q., Li, S., Liang, S., Zhong, H., Liu, Z., Gao, Y., Zhao, H., et al. (2017). The gut microbiome in atherosclerotic cardiovascular disease. Nat. Commun. 8, 845.

41. Karlsson, F.H., Tremaroli, V., Nookaew, I., Bergström, G., Behre, C.J., Fagerberg, B., Nielsen, J., and Bäckhed, F. (2013). Gut metagenome in European women with normal, impaired and diabetic glucose control. Nature 498, 99–103.

42. Lewis, J.D., Chen, E.Z., Baldassano, R.N., Otley, A.R., Griffiths, A.M., Lee, D., Bittinger, K., Bailey, A., Friedman, E.S., Hoffmann, C., et al. (2015). Inflammation, antibiotics, and diet as environmental stressors of the gut microbiome in pediatric Crohn’s disease. Cell Host Microbe 18, 489–500.

43. Lloyd-Price, J., Arze, C., Ananthakrishnan, A.N., Schirmer, M., Avila-Pacheco, J., Poon, T.W., Andrews, E., Ajami, N.J., Bonham, K.S., Brislawn, C.J., et al. (2019). Multi-omics of the gut microbial ecosystem in inflammatory bowel diseases. Nature 569, 655–662.

44. Loomba, R., Seguritan, V., Li, W., Long, T., Klitgord, N., Bhatt, A., Dulai, P.S., Caussy, C., Bettencourt, R., Highlander, S.K., et al. (2017). Gut microbiome-based metagenomic signature for non-invasive detection of advanced fibrosis in human nonalcoholic fatty liver disease. Cell Metab. 25, 1054–1062. e5.

45. Mehta, R.S., Abu-Ali, G.S., Drew, D.A., Lloyd-Price, J., Subramanian, A., Lochhead, P., Joshi, A.D., Ivey, K.L., Khalili, H., Brown, G.T., et al. (2018). Stability of the human faecal microbiome in a cohort of adult men. Nat. Microbiol. 3, 347–355.

46. Qin, J., Li, Y., Cai, Z., Li, S., Zhu, J., Zhang, F., Liang, S., Zhang, W., Guan, Y., Shen, D., et al. (2012). A metagenome-wide association study of gut microbiota in type 2 diabetes. Nature 490, 55–60.

47. Qin, N., Yang, F., Li, A., Prifti, E., Chen, Y., Shao, L., Guo, J., Le Chatelier, E., Yao, J., Wu, L., et al. (2014). Alterations of the human gut microbiome in liver cirrhosis. Nature 513, 59–64.

48. Yang, K., Niu, J., Zuo, T., Sun, Y., Xu, Z., Tang, W., Liu, Q., Zhang, J., Ng, E.K., Wong, S.K., et al. (2021). Alterations in the gut virome in obesity and type 2 diabetes mellitus. Gastroenterology 161, 1257–1269. e13.

49. Zhang, X., Zhang, D., Jia, H., Feng, Q., Wang, D., Liang, D., Wu, X., Li, J., Tang, L., Li, Y., et al. (2015). The oral and gut microbiomes are perturbed in rheumatoid arthritis and partly normalized after treatment. Nat. Med. 21, 895–905.

50. Li, J., Zhao, F., Wang, Y., Chen, J., Tao, J., Tian, G., Wu, S., Liu, W., Cui, Q., Geng, B., et al. (2017). Gut microbiota dysbiosis contributes to the development of hypertension. Microbiome 5, 1–19.

51. Li, Z., Zhou, J., Liang, H., Ye, L., Lan, L., Lu, F., Wang, Q., Lei, T., Yang, X., and Cui, P. (2022). Differences in alpha diversity of gut microbiota in neurological diseases. Front. Neurosci. 16, 879318.

52. Gong, C.H., Kendig, H., and He, X. (2016). Factors predicting health services use among older people in China: an analysis of the China health and retirement longitudinal study 2013. BMC Health Serv. Res. 16, 63.

53. Gacesa, R., Kurilshikov, A., Vich Vila, A., Sinha, T., Klaassen, M., Bolte, L., Andreu-Sánchez, S., Chen, L., Collij, V., Hu, S., et al. (2022). Environmental factors shaping the gut microbiome in a Dutch population. Nature 604, 732–739.

54. Gupta, V.K., Kim, M., Bakshi, U., Cunningham, K.Y., Davis III, J.M., Lazaridis, K.N., Nelson, H., Chia, N., and Sung, J. (2020). A predictive index for health status using species-level gut microbiome profiling. Nat. Commun. 11, 4635.

55. Aron-Wisnewsky, J., Vigliotti, C., Witjes, J., Le, P., Holleboom, A.G., Verheij, J., Nieuwdorp, M., and Clément, K. (2020). Gut microbiota and human NAFLD: disentangling microbial signatures from metabolic disorders. Nat. Rev. Gastroenterol. Hepatol. 17, 279–297.

56. Zeisel, S.H., and Warrier, M. (2017). Trimethylamine N-oxide, the microbiome, and heart and kidney disease. Annu. Rev. Nutr. 37, 157–181.

57. Falony, G., Vieira-Silva, S., and Raes, J. (2015). Microbiology meets big data: the case of gut microbiota–derived trimethylamine. Annu. Rev. Microbiol. 69, 305–321.

58. Cai, Y.-Y., Huang, F.-Q., Lao, X., Lu, Y., Gao, X., Alolga, R.N., Yin, K., Zhou, X., Wang, Y., Liu, B., et al. (2022). Integrated metagenomics identifies a crucial role for trimethylamine-producing *Lachnoclostridium* in promoting atherosclerosis. NPJ Biofilms Microbiomes 8, 11.

59. Dalla Via, A., Gargari, G., Taverniti, V., Rondini, G., Velardi, I., Gambaro, V., Visconti, G.L., De Vitis, V., Gardana, C., Ragg, E., et al. (2019). Urinary TMAO levels are associated with the taxonomic composition of the gut microbiota and with the choline TMA-lyase gene (*cutC*) harbored by Enterobacteriaceae. Nutrients 12, 62.

60. Estrada, A., Garber, P.A., Rylands, A.B., Roos, C., Fernandez-Duque, E., Di Fiore, A., Nekaris, K.A.-I., Nijman, V., Heymann, E.W., Lambert, J.E., et al. (2017). Impending extinction crisis of the world’s primates: Why primates matter. Sci. Adv. 3, e1600946.

61. Grieneisen, L.E., Charpentier, M.J., Alberts, S.C., Blekhman, R., Bradburd, G., Tung, J., and Archie, E.A. (2019). Genes, geology and germs: gut microbiota across a primate hybrid zone are explained by site soil properties, not host species. Proc. R. Soc. B 286, 20190431.

62. Ben-Dor, M., Sirtoli, R., and Barkai, R. (2021). The evolution of the human trophic level during the Pleistocene. Am. J. Phys. Anthropol. 175, 27–56.

63. Groussin, M., Poyet, M., Sistiaga, A., Kearney, S.M., Moniz, K., Noel, M., Hooker, J., Gibbons, S.M., Segurel, L., Froment, A., et al. (2021). Elevated rates of horizontal gene transfer in the industrialized human microbiome. Cell 184, 2053–2067. e18.

64. Zhao, S., Lieberman, T.D., Poyet, M., Kauffman, K.M., Gibbons, S.M., Groussin, M., Xavier, R.J., and Alm, E.J. (2019). Adaptive evolution within gut microbiomes of healthy people. Cell Host Microbe 25, 656–667. e8.

65. De Filippis, F., Pasolli, E., Tett, A., Tarallo, S., Naccarati, A., De Angelis, M., Neviani, E., Cocolin, L., Gobbetti, M., Segata, N., and Ercolini D. (2019). Distinct genetic and functional traits of human intestinal *Prevotella copri* strains are associated with different habitual diets. Cell Host Microbe 25, 444–453. e3.

66. Hosokawa, T., Kikuchi, Y., Nikoh, N., Shimada, M., and Fukatsu, T. (2006). Strict host-symbiont cospeciation and reductive genome evolution in insect gut bacteria. PLoS Biol. 4, e337.

67. McCutcheon, J.P., and Moran, N.A. (2012). Extreme genome reduction in symbiotic bacteria. Nat. Rev. Microbiol. 10, 13–26.

68. Beasley, D.E., Koltz, A.M., Lambert, J.E., Fierer, N., and Dunn, R.R. (2015). The evolution of stomach acidity and its relevance to the human microbiome. PLoS One 10, e0134116.

69. Qiao, Y., Sun, J., Ding, Y., Le, G., and Shi, Y. (2013). Alterations of the gut microbiota in high-fat diet mice is strongly linked to oxidative stress. Appl. Microbiol. Biotechnol. 97, 1689–1697.

70. Vaccaro, A., Dor, Y.K., Nambara, K., Pollina, E.A., Lin, C., Greenberg, M.E., and Rogulja, D. (2020). Sleep loss can cause death through accumulation of reactive oxygen species in the gut. Cell 181, 1307–1328. e1315.

71. Sistiaga, A., Wrangham, R., Rothman, J.M., and Summons, R.E. (2015). New insights into the evolution of the human diet from faecal biomarker analysis in wild chimpanzee and gorilla faeces. PLoS One 10, e0128931.

72. Hecker, N., Sharma, V., and Hiller, M. (2019). Convergent gene losses illuminate metabolic and physiological changes in herbivores and carnivores. Proc. Natl. Acad. Sci. USA 116, 3036–3041.

73. Johnson, C.A., Smith, G.P., Yule, K., Davidowitz, G., Bronstein, J.L., and Ferrière, R. (2021). Coevolutionary transitions from antagonism to mutualism explained by the co-opted antagonist hypothesis. Nat. Commun. 12, 2867.

74. Blaser, M.J. (2017). The theory of disappearing microbiota and the epidemics of chronic diseases. Nat. Rev. Immunol. 17, 461–463.

75. Schirmer, M., Garner, A., Vlamakis, H., and Xavier, R.J. (2019). Microbial genes and pathways in inflammatory bowel disease. Nat. Rev. Microbiol. 17, 497–511.

76. Khan Mirzaei, M., and Deng, L. (2021). Sustainable microbiome: a symphony orchestrated by synthetic phages. Microb. Biotechnol. 14, 45–50.

77. Degnan, P.H., Barry, N.A., Mok, K.C., Taga, M.E., and Goodman, A.L. (2014). Human gut microbes use multiple transporters to distinguish vitamin B12 analogs and compete in the gut. Cell Host Microbe 15, 47–57.

78. Bentkowski, P., Van Oosterhout, C., and Mock, T. (2015). A model of genome size evolution for prokaryotes in stable and fluctuating environments. Genome Biol. Evol. 7, 2344–2351.

79. Delannoy-Bruno, O., Desai, C., Raman, A.S., Chen, R.Y., Hibberd, M.C., Cheng, J., Han, N., Castillo, J.J., Couture, G., Lebrilla, C.B., et al. (2021). Evaluating microbiome-directed fibre snacks in gnotobiotic mice and humans. Nature 595, 91–95.

80. Zhang, C., Björkman, A., Cai, K., Liu, G., Wang, C., Li, Y., Xia, H., Sun, L., Kristiansen, K., Wang, J., et al. (2018). Impact of a 3-months vegetarian diet on the gut microbiota and immune repertoire. Front. Immunol. 9, 908.

81. Youngblut, N.D., Reischer, G.H., Walters, W., Schuster, N., Walzer, C., Stalder, G., Ley, R.E., and Farnleitner, A.H. (2019). Host diet and evolutionary history explain different aspects of gut microbiome diversity among vertebrate clades. Nat. Commun. 10, 2200.

82. Bolger, A.M., Lohse, M., and Usadel, B. (2014). Trimmomatic: a flexible trimmer for Illumina sequence data. Bioinformatics 30, 2114–2120.

83. Nurk, S., Meleshko, D., Korobeynikov, A., and Pevzner, P.A. (2017). metaSPAdes: a new versatile metagenomic assembler. Genome Res. 27, 824–834.

84. Li, D., Liu, C.-M., Luo, R., Sadakane, K., and Lam, T.-W. (2015). MEGAHIT: an ultra-fast single-node solution for large and complex metagenomics assembly via succinct de Bruijn graph. Bioinformatics 31, 1674–1676.

85. Uritskiy, G.V., DiRuggiero, J., and Taylor, J. (2018). MetaWRAP—a flexible pipeline for genome-resolved metagenomic data analysis. Microbiome 6, 158.

86. Wu, Y.-W., Simmons, B.A., and Singer, S.W. (2016). MaxBin 2.0: an automated binning algorithm to recover genomes from multiple metagenomic datasets. Bioinformatics 32, 605–607.

87. Kang, D.D., Froula, J., Egan, R., and Wang, Z. (2015). MetaBAT, an efficient tool for accurately reconstructing single genomes from complex microbial communities. PeerJ 3, e1165.

88. Parks, D.H., Imelfort, M., Skennerton, C.T., Hugenholtz, P., and Tyson, G.W. (2015). CheckM: assessing the quality of microbial genomes recovered from isolates, single cells, and metagenomes. Genome Res. 25, 1043–1055.

89. Olm, M.R., Brown, C.T., Brooks, B., and Banfield, J.F. (2017). dRep: a tool for fast and accurate genomic comparisons that enables improved genome recovery from metagenomes through de-replication. ISME J. 11, 2864–2868.

90. Langmead, B., and Salzberg, S.L. (2012). Fast gapped-read alignment with Bowtie 2. Nat. Methods 9, 357–359.

91. Chaumeil, P.-A., Mussig, A.J., Hugenholtz, P., and Parks, D.H. (2020). GTDB-Tk: a toolkit to classify genomes with the Genome Taxonomy Database. Bioinformatics 36, 1925–1927.

92. Katoh, K., and Standley, D.M. (2013). MAFFT multiple sequence alignment software version 7: improvements in performance and usability. Mol. Biol. Evol. 30, 772–780.

93. Price, M.N., Dehal, P.S., and Arkin, A.P. (2010). FastTree 2–approximately maximum-likelihood trees for large alignments. PLoS One 5, e9490.

94. Letunic, I., and Bork, P. (2019). Interactive Tree Of Life (iTOL) v4: recent updates and new developments. Nucleic Acids Res. 47, (W1), W256–W259.

95. Hyatt, D., Chen, G.-L., LoCascio, P.F., Land, M.L., Larimer, F.W., and Hauser, L.J. (2010). Prodigal: prokaryotic gene recognition and translation initiation site identification. BMC Bioinformatics 11, 119.

96. Cantalapiedra, C.P., Hernández-Plaza, A., Letunic, I., Bork, P., and Huerta-Cepas, J. (2021). eggNOG-mapper v2: functional annotation, orthology assignments, and domain prediction at the metagenomic scale. Mol. Biol. Evol. 38, 5825–5829.

97. Huerta-Cepas, J., Szklarczyk, D., Heller, D., Hernández-Plaza, A., Forslund, S.K., Cook, H., Mende, D.R., Letunic, I., Rattei, T., and Jensen, L.J. (2019). eggNOG 5.0: a hierarchical, functionally and phylogenetically annotated orthology resource based on 5090 organisms and 2502 viruses. Nucleic Acids Res. 47, D309–D314.

98. Zhang, H., Yohe, T., Huang, L., Entwistle, S., Wu, P., Yang, Z., Busk, P.K., Xu, Y., and Yin, Y. (2018). dbCAN2: a meta server for automated carbohydrate-active enzyme annotation. Nucleic Acids Res. 46, W95–W101.

99. Camacho, C., Coulouris, G., Avagyan, V., Ma, N., Papadopoulos, J., Bealer, K., and Madden, T.L. (2009). BLAST+: architecture and applications. BMC Bioinformatics 10, 421.

100. Arango-Argoty, G., Garner, E., Pruden, A., Heath, L.S., Vikesland, P., and Zhang, L. (2018). DeepARG: a deep learning approach for predicting antibiotic resistance genes from metagenomic data. Microbiome 6, 23.

101. Stamatakis, A. (2014). RAxML version 8: a tool for phylogenetic analysis and post-analysis of large phylogenies. Bioinformatics 30, 1312–1313.

102. Fu, L., Niu, B., Zhu, Z., Wu, S., and Li, W. (2012). CD-HIT: accelerated for clustering the next-generation sequencing data. Bioinformatics 28, 3150–3152.

103. Buchfink, B., Xie, C., and Huson, D.H. (2015). Fast and sensitive protein alignment using DIAMOND. Nat. Methods 12, 59–60.

104. Beghini, F., McIver, L.J., Blanco-Míguez, A., Dubois, L., Asnicar, F., Maharjan, S., Mailyan, A., Manghi, P., Scholz, M., Thomas, A.M., et al. (2021). Integrating taxonomic, functional, and strain-level profiling of diverse microbial communities with bioBakery 3. eLife 10, e65088.

105. Pedregosa, F., Varoquaux, G., Gramfort, A., Michel, V., Thirion, B., Grisel, O., Blondel, M., Prettenhofer, P., Weiss, R., and Dubourg, V. (2011). Scikit-learn: machine learning in Python. J. Mach. Learn. Res. 12, 2825–2830.

106. Watts, S.C., Ritchie, S.C., Inouye, M., and Holt, K.E. (2019). FastSpar: rapid and scalable correlation estimation for compositional data. Bioinformatics 35, 1064–1066.

107. Smoot, M.E., Ono, K., Ruscheinski, J., Wang, P.-L., and Ideker, T. (2011). Cytoscape 2.8: new features for data integration and network visualization. Bioinformatics 27, 431–432.

108. Oksanen, J., Simpson, G., Blanchet, F., Kindt, R., Legendre, P., Minchin, P., O’Hara, R., Solymos, P., Stevens, M., Szoecs, E., et al. (2022). vegan: Community Ecology Package. R package version 2.6-2.

