## Supplemental Text and Figures for "Opposing implications of co-evolutionary lineages and traits of gut microbiome on human health status"

**Supplementary figures and legends**

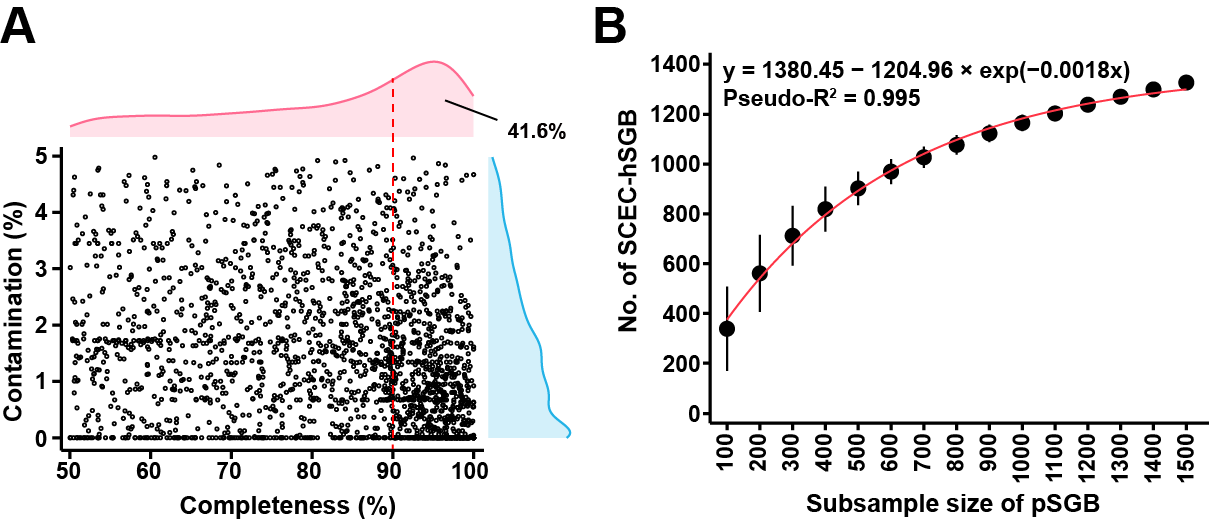

**Figure S1, A) genome quality of 2,036 SGBs recovered in this study** **and B) rarefaction curves of the number of SCEC-hSGB versus pSGB subsample size (mean ± s.d.), related to Figure 1.**

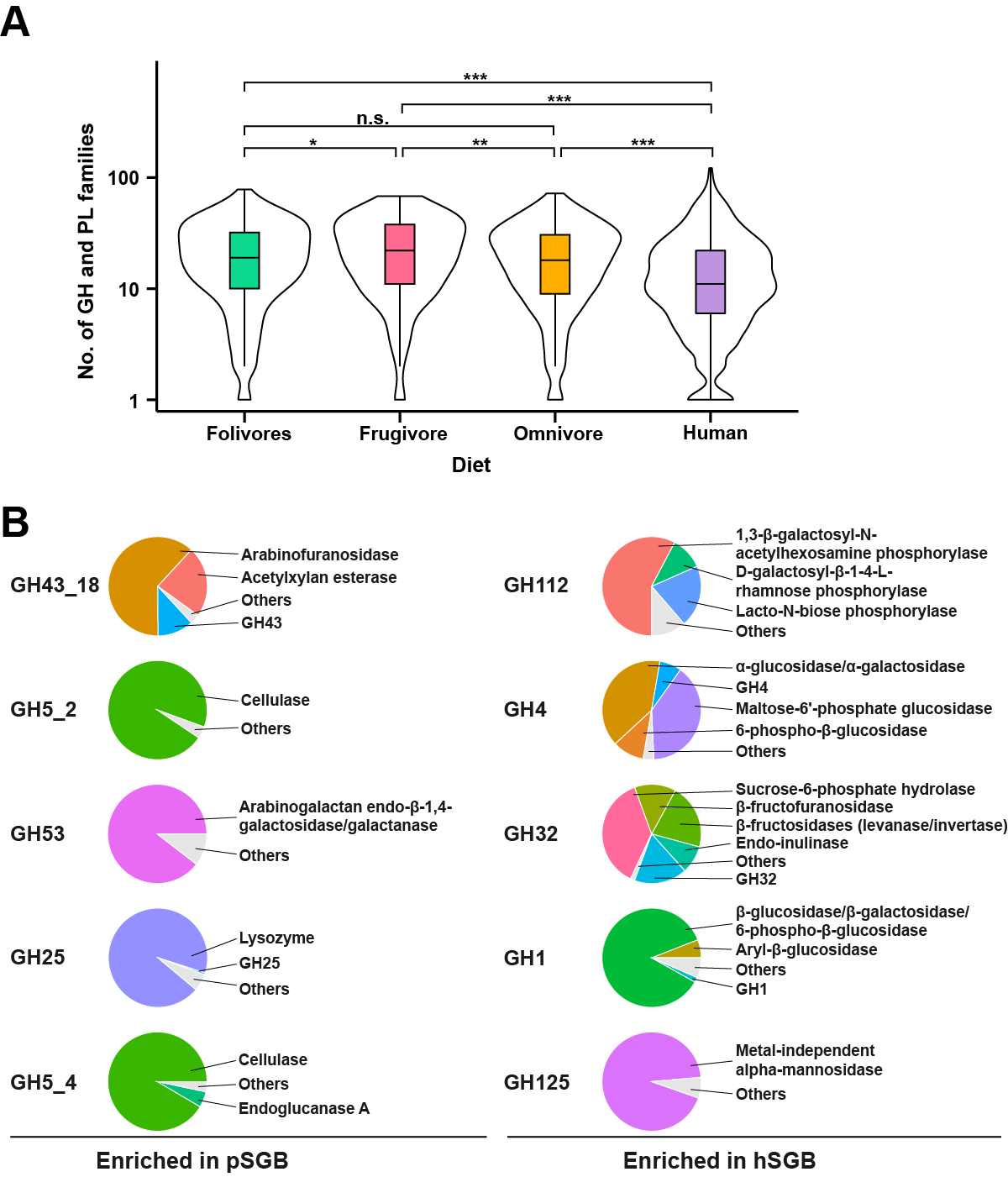

**Figure S2, CAZy profiling of pSGB and hSGB, related to Figure 2.**

**A)** The number of GH and PL families in SGBs from different diet group. Two-tailed Mann-Whitney U-test. *, *P*_adj_ < 0.05; **, *P*_adj_ < 0.01; ***, *P*_adj_ < 0.001; n.s., not significant. **B)** The substrates categories of the top 5 enriched CAZymes in pSGBs or hSGBs.

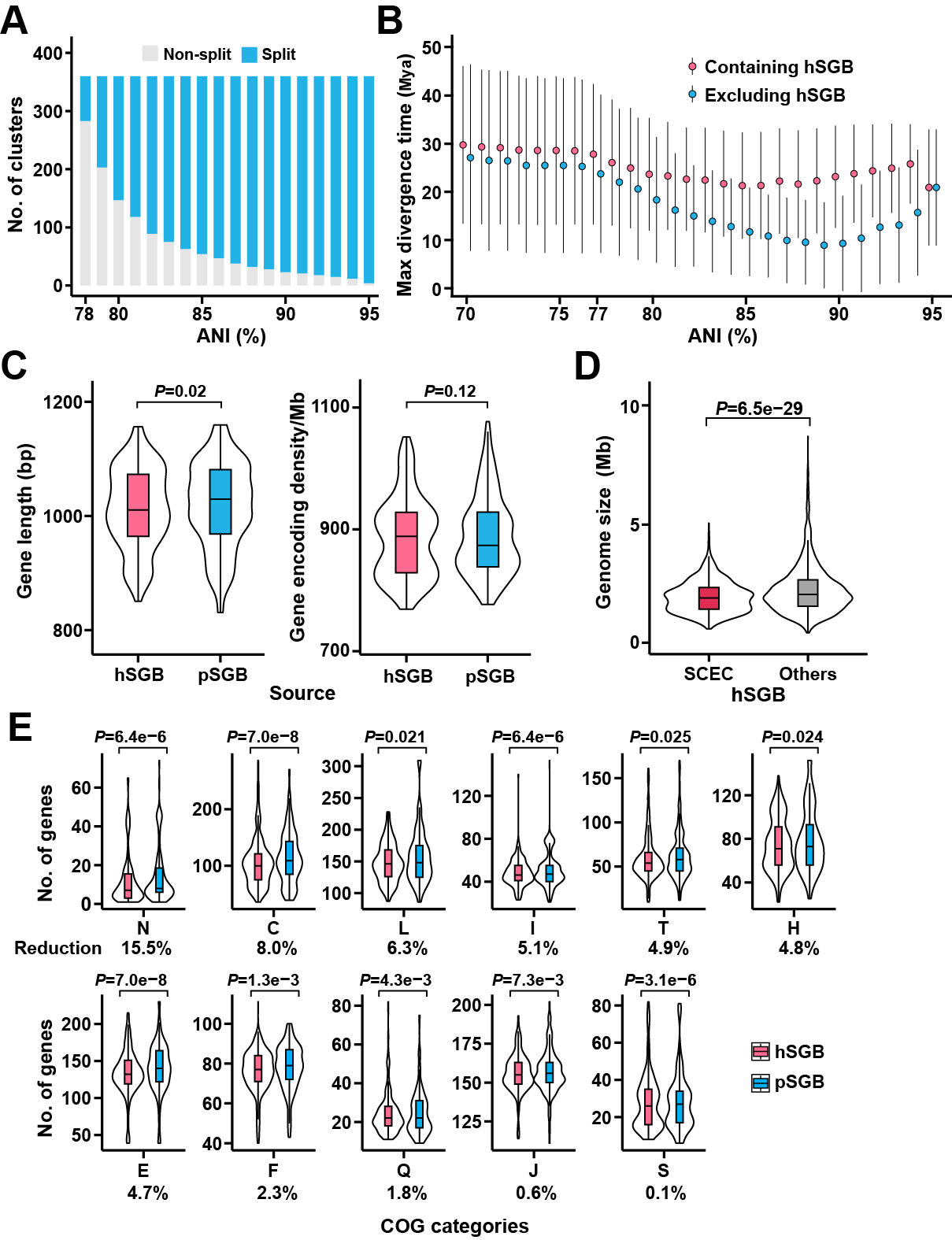

**Figure S3, defining and characterization of co-evolutionary SGB clusters, related to Figure 3. A)** The split ratio of ANI-77% non-singleton clusters at subsequent thresholds. **B)** The maximum divergence time of hosts corresponding to the SGBs within the cluster. Each point represents the average maximum divergence time of all clusters under that threshold. Error bars indicate the standard deviations. **C)** Comparison of gene length and coding density between pSGBs and hSGBs within SCEC. Only SGBs with completeness >95% and complete ORFs were considered. Paired two-sided student’s *t*-test. **D)** Comparison of genome size between SCEC-hSGBs and other hSGBs. Two-sided Student’s *t*-test. **E)** The enriched COG categories of SCEC-pSGBs compared to SCEC-hSGBs. Only SGBs with completeness >90% were considered. The reduction of hSGB compared to pSGB in each cog category is listed below the plot. Paired two-sided Student’s *t*-test with FDR correction.

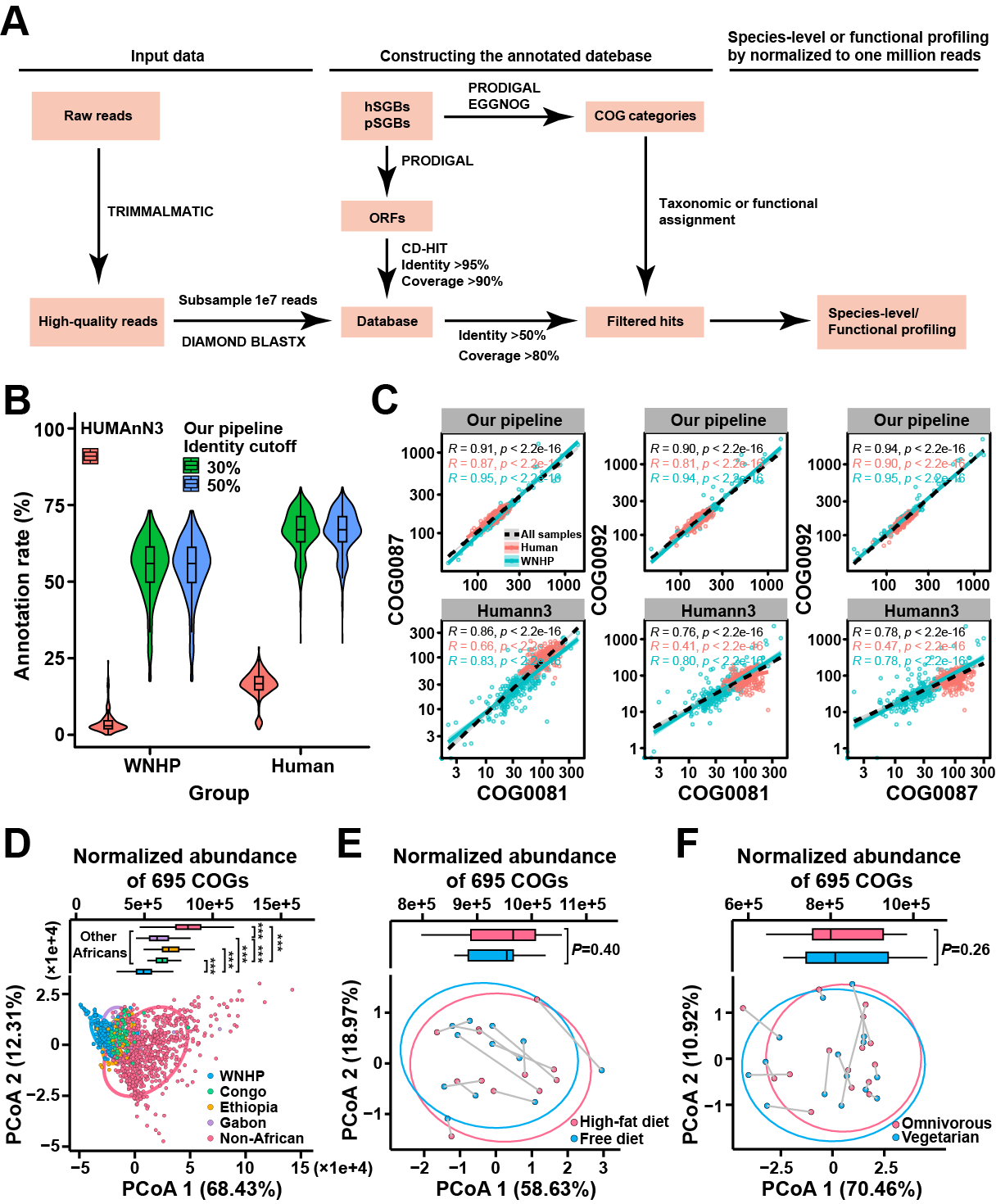

**Figure S4, the quantification profiling pipeline in metagenomes and characterization co-evolutionary traits in different metagenome groups, related to Figure 4. A)** The workflow of the profiling pipeline. We built a protein database containing ORFs from pSGBs and hSGBs. In the search stage, quality-filtered reads were aligned against the custom database. In the annotation stage, we use the total number of hits annotated to ORFs within the same ortholog as the abundance of that COG. Comparison of **B)** the annotation rate and **C)** Spearman's ρ among three conserved proteins using our pipeline and HUMAnN3 on the metagenome of WHNP and human. **D)** The distribution of 695 COGs in WNHP and different human populations. PCoA based on the 695 COGs, Euclidean distance. Ellipses cover 90% of the metagenome for each group. Two-sided Student’s *t*-test with FDR correction. *, *P*_adj_ < 0.05; **, *P*_adj_ < 0.01; ***, *P*_adj_ < 0.001; n.s., not significant. **E, F)** The effect of short-term dietary intervention on the abundance of 695 COGs. Ellipses cover 90% of the metagenome for each group. Paired two-sided Student’s *t*-test. **E)** Dietary intervention (from free diet to diet with high in saturated fats and low in fruits and vegetables, ~1 week) involving 12 overweight volunteers. **F)** A three-month dietary intervention (from omnivorous diet to vegetarian diet for three months) involving 15 volunteers.

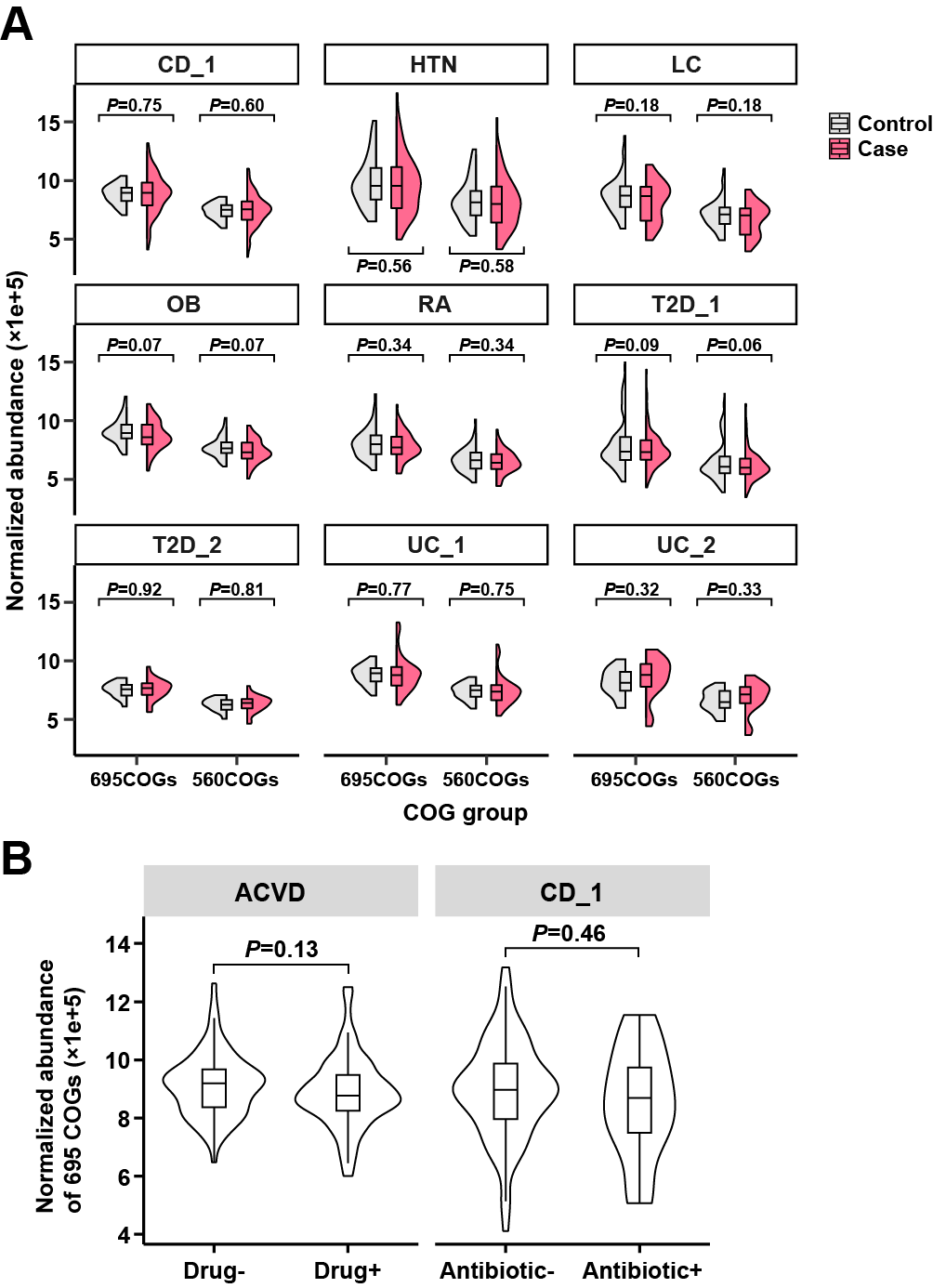

**Figure S5,** **the associations of the traits COGs with disease, related to Figure 5. A)** The abundance difference of 695 COGs and 560 COGs between case and control groups. **B)** The effect of drug and antibiotic usage on the 695 COGs abundance. Two-sided Student’s *t*-test.

HTN, hypertension; LC, liver cirrhosis; CD Crohn’s disease; OB, obesity; RA, rheumatoid arthritis; T2D, type 2 diabetes; UC, ulcerative colitis.

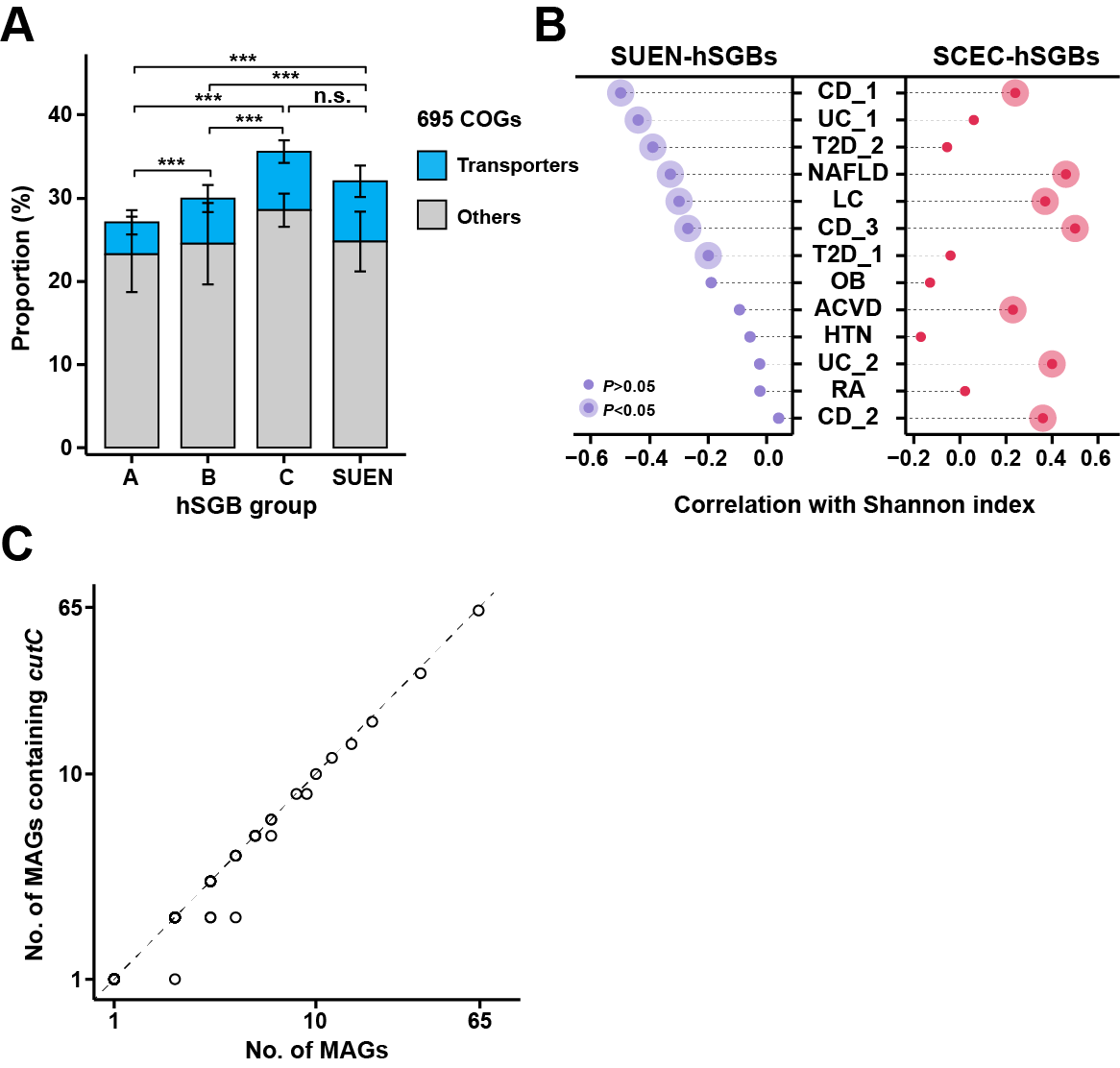

**Figure S6, characterization of 695 COGs and indications for host health of SCEC- and SUNE-hSGBs, related to Figure 6. A)** The proportion of 695 COGs and the transporter in hSGBs. Mean ± s.d. Two-sided Mann-Whitney U-test with FDR correction for the proportion of transporters. *, *P*_adj_ < 0.05; **, *P*_adj_ < 0.01; ***, *P*_adj_ < 0.001; n.s., not significant. **B)** Spearman's correlation of the relative abundance of SUEN- and SCEC-hSGBs in metagenome and its Shannon index. The SCEC- or SUEN-hSGBs were taken into account when calculating the Shannon index. **C)** The intra-species variation of encoding *CutC*. Only MAGs with completeness >90% were considered.

ACVD, atherosclerotic cardiovascular disease; NAFLD, nonalcoholic fatty liver disease; HTN, hypertension; LC, liver cirrhosis; CD, Crohn’s disease; OB, obesity; RA, rheumatoid arthritis; T2D, type 2 diabetes; UC, ulcerative colitis.
